# Partitioning of nighttime transpiration and stem water refilling using VPD and dendrometer data: insights into baselining and nighttime sap flux interpretation

**DOI:** 10.1101/2025.08.25.672109

**Authors:** Mianzhi Wang, Matti Räsänen, Teemu Hölttä

**Author notes:** **Author for correspondence:** Mianzhi Wang, Latokartanonkaari 7, 00014 University of Helsinki, Finland.

## Abstract

The baseline for thermal dissipation probes is a reference for simultaneous cessation of both nighttime transpiration and refilling of trees internal water stores. Finding this moment is challenging as the baseline is dynamically changing, and the complete sap flux cessation may not even occur. We proposed a novel framework using VPD and dendrometer measurements to partition transpiration and refilling from nighttime sap flux density, and introduced a night-by-night baselining scheme that does not rely on identifying the zero-flow moment. One conventional baselining method using the maximum probe voltage (or temperature) difference substantially underestimated sap flux density by 204.56% for nighttime, 28.30% for daytime, and 45.20% for full-day flux in *P. sylvestris*, and by 344.77%, 45.68%, and 54.76%, respectively, in *A. glutinosa*. Baseline calibrations using commonly available variables are thus provided. Evaluation through gas exchange and evapotranspiration data confirmed the accuracy of partitioning between nighttime transpiration and refilling, demonstrating that nighttime transpiration can drive measurable nighttime evapotranspiration responses. Nightly variations in water allocation within stem tissue add complexity to stem swelling. Seasonally, sap flux density showed an initial increase in both transpiration and refilling, followed by a decline, with transpiration dominating until mid-summer before refilling gradually took precedence.

**Highlights:**

- Partitioned nighttime transpiration & stem refilling using VPD & dendrometers.
- Tracked seasonal shifts in transpiration & refilling ratios to nighttime sap flux.
- Independently evaluated model with eddy covariance & leaf gas exchange measurements.
- Proposed a new high-frequency baselining strategy for thermal dissipation probes.
- Revealed & corrected errors in sap flux density baselining methods.

## Introduction

Many studies have shown that trees have nighttime sap flux for transpiration and stem water refilling (Chowdhury et al., 2022; Wang et al., 2012; Zeppel et al., 2012; Zhao et al., 2019, 2022). Nighttime transpiration includes water loss through partly opened stomata and residual water loss through leaf cuticle, as well as water loss through the bark of stem and branches (Brito et al., 2018; Lintunen et al., 2021; Novick et al., 2009; Zeppel et al., 2014). When less water is transpired at night than is taken up from the soil, the stem water is refilled. This process replenishes the stem’s water lost during the day, maintains adequate stem turgor, and supports turgor-dependent growth (Chan et al., 2016; Peters et al., 2023; Zweifel et al., 2021). This replenishment leads to stem swelling, which can be detected using dendrometers (Zeppel et al., 2010). Changes in stem water content were closely associated with changes in stem diameter, and in particular changes in stem symplastic water content could be monitored by the dendrometer, while changes in apoplastic water content were not as effectively monitored (Lintunen et al., 2017). Transpiration and refilling both lead to sap flux density, which can be monitored using various methods such as Granier-type thermal dissipation probes (TDP), the heat field deformation method (HFD), and the heat ratio method (HRM) (Fuchs et al., 2017). Thus, growing efforts focus on integrating sap flow sensors and dendrometers to deepen understanding of tree stem water dynamics (De Pauw et al., 2008; Oliva Carrasco et al., 2015; Steppe et al., 2006). Although standard TDP and datalogger system require calibration to accurately quantify sap flow (Fuchs et al., 2017; Peters et al., 2018), they are widely used due to their simplicity, cost-effectiveness, and reliability of capturing sap flow relative changes (Flo et al., 2019; Poyatos et al., 2016). Nonetheless, two challenges have hindered the evaluation of nighttime sap flux for TDP users: (1) determining the baseline, i.e. the points in time where sap flux density is zero, and (2) partitioning between nighttime transpiration and refilling (Fisher et al., 2007; Oishi et al., 2016).

As the two fundamental processes of nighttime sap flux activities are transpiration and refilling (Fisher et al., 2007), the cessation of sap flux indicates that both processes cease simultaneously. Previous studies have commonly defined the cessation of sap flux based on low evaporative demand and the stability of ΔT or ΔV, an approach that has proven effective across various tree species and environments (Di et al., 2019; Oishi et al., 2016; Yu et al., 2018; Zeppel et al., 2010). However, in practice, such moments are challenging to identify and may not even occur throughout the growing season in certain environments for some species. Moreover, although low VPD is well used to determine cessation of transpiration, few quantitative metrics have been used to determine cessation of refilling. In addition, the baseline may change with stem water content (Hölttä et al. 2015) and the surrounding environment (Hölttä et al., 2015; Oishi et al., 2016), suggesting that methods including a multi-day moving window are also not ideal as it may obscure these important variation in the baseline (Lu et al., 2004). Thus, continuous, and accurate monitoring of nighttime sap flux demands frequent baselining and the consideration of both transpiration and refilling, entailing the need to partition between the two processes.

However, there has not been a flawless scheme for partitioning between transpiration and refilling in previous studies. The time-separation approach is a classic concept proposed to directly partition refilling and transpiration primarily based on an modelling the refiling part of the sap flux as an exponential decay function (Fisher et al., 2007). While this approach has been widely adopted (Alvarado-Barrientos et al., 2015; Yu et al., 2018; Zhao et al., 2019), it lacks a strong theoretical and practical foundations. Currently, purely model-based partitioning methods have limited evaluation through direct measurements (Zhao et al., 2022), while those based on gas exchange measurements are difficult to monitor over time (Wang et al., 2012).

Here, we provide a new approach based on a combination of continuous VPD and dendrometer measurements to effectively partition between transpiration and refilling with high temporal resolution. The nighttime sap flux density consists of two components: one supporting transpiration and the other refilling of stem tissues which were depleted during the daytime and water taken for cambial growth. The portion of flux used for transpiration is primarily driven by VPD, while the flux used for refilling is reflected in stem radial swelling. Based on this principle, our approach establishes a link between sap flux density, VPD, and stem diameter. In addition, we propose a new baselining strategy that incorporates both processes. In boreal regions, where the summer nights are short and sap flux density may not decrease to zero during night, two widely distributed tree species, Scots pine (*P. sylvestris*) and black alder (*A. glutinosa*) (Brichta et al., 2023; Claessens et al., 2010), were used for our study. Our objectives were to: (1) introduce and evaluate a method for partitioning between nighttime sap flux used for transpiration and refilling, along with the associated effective baselining strategy, and (2) illustrate seasonal temporal patterns of transpiration and refilling.

## Materials and Methods Site

Two Scots pines (*P. sylvestris*) were measured in an even-aged homogeneous ~60 year-old Scots pine stand at SMEAR □ site in Hyytiälä, Finland (61°50′50″N, 24°17′41″E, 180 m a.s.l.) (Vesala et al., 1998). The previous stand was felled in 1961, followed by prescribed burning, after which Scots pine seeds were sown. The measured trees during the measurement year averaged approximately 18 m in height, with an average breast height diameter of 20 cm. The soil type is acidic glacial till with low bulk density (Gillespie et al., 2024; Lintunen et al., 2020). The measurements were conducted over the period from 2015 to 2017, with annual precipitation recorded at 756.59 mm and an average air temperature of +4.81□.

Two black alders (*A. glutinosa*) were measured in city of Helsinki, Finland (60°13′36″N, 25°1′41″E, 5 m a.s.l.) in 2011. These trees were planted along a street in 2002, spaced approximately 4-5 meters apart. No additional irrigation was provided beyond natural rainfall. The soil contains pre-mixed structural soil installed as planting pockets separated by compacted gravel (Havu et al., 2022; Lintunen et al., 2020). The average tree height and breast height diameter in 2010 were 11 m and 15 cm, respectively (Lintunen et al., 2020). In 2011, the annual precipitation was 733.57 mm, and average air temperature was +7.15□.

### Measurements

Sap flux density was measured every minute using TDP (Granier, 1985). Custom-made 4 cm long probes were inserted into northern side of the stem at a height of 10-12 m in *P. sylvestris* and 0.5-1 m in *A. glutinosa*, and were covered with brass sleeves and thermal paste. The voltage difference (ΔV) between the two probes was recorded in place of ΔT, as both yield equivalent sap flux density calculations due to unit cancellation in the formula. The probes were removed and reinstalled annually. The probe needles did not reach heartwood (Hölttä et al., 2010). Radial stem variations were measured with point-dendrometers (LVDT; model AX/5.0/S; Solartron Inc., West Sussex, UK). The measurement setup exhibited negligible thermal expansion, as the expansion of both the frame and wood nearly canceled each other out (Belder, n.d.; Sevanto et al., 2005) Additionally, nighttime temperatures at the study site remained relatively stable throughout the study period, with an average variation of 3.47°C in Hyytiälä and 3.29°C in Helsinki and the amplitude of the change in stem diameter during the nighttime being very similar as in the daytime, thus minimizing the potential impact of temperature on thermal expansion.

Air temperature and photosynthetic active radiation (PAR) were continuously measured at a height of 16 m at the SMEAR II station and 8 m at the urban site. When PAR fell below 10 µmol m□^2^ s□^1^ and 40 µmol m□^2^ s□^1^ (day/night threshold) in Hyytiälä and Helsinki, respectively, it was considered night. The Number of Night (NOY) and Number of Day (DOY) were determined based on natural sunrise and sunset. DOY 1 was defined as the period from sunrise to sunset on January 1^st^ when PAR exceeded the day/night threshold, while NOY 1 was the period from sunset on January 1^st^ to sunrise on January 2^nd^ when PAR fell below the threshold, and so forth.

The SWC, precipitation, daytime and nighttime VPD of the two sites during the study year are shown in Fig. S1. At the SMEAR II station, SWC was measured at depths ranging from 2 to 61 cm, whereas at the urban site it was measured at a depth of 10 cm using a Campbell TDR100 sensor. At SMEAR II, relative air humidity (Rotronic MP102H) and air temperature (Pt100) were measured at a height of 16.8 m, while precipitation (Vaisala FD12P weather sensor) was recorded at 18 m. For the urban site, relative air humidity was obtained from a nearby Vaisala automated weather station located approximately 400 m away, and precipitation data were acquired from the SMEAR III urban measurement station, situated 4 km from the site (Järvi et al., 2009). The vapor pressure deficit (VPD) was calculated from measurements of relative humidity (RH) and air temperature.

Eddy covariance measurements of evapotranspiration (ET) were obtained from the Hyytiälä SMEAR II station at a height of 24 meters (Junninen et al., 2009). The ET values were discarded during rainy periods, when the friction velocity u* was below 0.38 m s^−1^ and when flux quality value was 2 (Mauder and Foken, 2015).

Leaf gas exchange measurements of transpiration rate (T_measure_) were conducted on one Scots pine in 2015 and another in 2016 and 2017. Measurement chambers were fitted onto shoots, each accommodating needles from one age class. The chambers were equipped with a pneumatically operated system that opened and closed approximately every 20 minutes, enabling automated and long-term monitoring of H_2_O fluxes (Kolari et al., 2014). To prevent new growth within the chambers, terminal buds were removed before installing the chambers. Following the gas exchange measurements, the dimensions of needles on each shoot were recorded, and their surface area was calculated using the equation from Tirén (1927), with the results divided by three to determine the projected area.

### Theory

Since there were very few nights in our dataset (only 7 nights out of four observation years) that satisfy the traditional “stable ΔV” criterion (standard deviation of ΔV is less than 0.5% of its mean during the two hours with the highest ΔV values, and VPD is less than 0.05 kPa), the nighttime sap flux density (Jn_0_, cm s^-1^) was initially calculated according to Grainer, 1985:

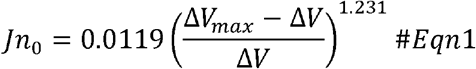

where ΔV is the voltage difference between the heating and reference probe, and ΔV_max_ is the maximum value of ΔV for each night. The “dTmax baseline” function of the Baseliner program was used to calculate this process (Oishi et al., 2016), which uses the ΔV_max_ as anchor points and can be seen as using the ΔV_max_ of each night as a baseline.

Nighttime sap flux density should be explained by both transpiration and refilling:

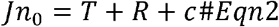

where Jn_0_, T, R, and c represent nighttime sap flux density (cm s^-1^), transpiration flux density (cm s^-1^), refilling flux density (cm s^-1^), and a constant, respectively. As the baseline (ΔV_max_) used to calculate Jn_0_ in Eqn1 may not always represent the true baseline, an intercept ‘c’ was introduced in the model to correct potential systematic shifts between Jn_0_ and the actual sap flux density. This adjustment ensures that the calculations of T and R remain equivalent to those obtained using the correct baseline. T is driven by VPD and is linearly related to VPD at a given canopy conductance (g_c_, sometimes g_c_ is referred to as “tree conductance” (e.g. Arneth et al. 1996, Whitehead et al. 1996)). R leads to stem swelling and is linearly correlated with the rate of stem swelling at a given stem water status. Based on these relationships, transpiration and refilling were modelled as follows:

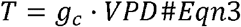

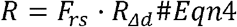

where VPD, R_Δd_, g_c_, and F_rs_ represent vapour pressure-deficit (kPa), rate of change in stem diameter obtained from the dendrometer measurements (μm s^-1^), canopy conductance (transpiration flux density per unit of VPD induced in transpiration process, cm s^-1^ kPa^-1^), and refilling swelling factor (refilling sap flux density per unit of R_Δd_ induced in the refilling process, unitless), respectively.

To obtain g_c_, F_rs_, and c, the multiple linear regression model (combination of Eqn2, Eqn3, and Eqn4) was applied on a night-by-night basis. It was assumed that g_c_, F_rs_, and c remained constant within each night, as nighttime duration is short, and the physiological conditions do not likely change significantly over a few hours. However, these parameters could vary from night to night. VPD data with a series of shifted timestamps was used to build model (Eqn2-4) to test potential the time lag of T response to VPD, and minimal time lag (within 5 minutes) was found and thus this time lag was not considered in further analysis (Fig. S2).

After modeling the data for all nights, we assessed model performance by setting specific criteria: the adjusted R^2^ (R_Adj_^2^) needed to exceed 0.8 to ensure a good fit, and both g_c_ and F_rs_ had to be significant (*P* < 0.05) and positive to confirm that the estimated parameters were statistically and biologically meaningful. Only models meeting these criteria were considered reliable and included in further analyses. The time series of g_c_ and F_rs_ are presented in Fig. S3. After obtaining the g_c_ and F_rs_ for each night, T and R were calculated according to Eqn3 and Eqn4.

Further, the sum of T and R is the modelled sap flux density (Jn_model_, cm s^-1^):

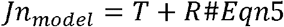

By combining Eqn2 and Eqn5, the intercept “c” accounts for the systematic difference between Jn_0_ and Jn_model_:

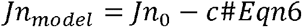

Each night, when ΔV = ΔV_max_, Jn_0_ equals to 0, while the actual Jn (Jn_model_ and the sap flux density calculated with correct baseline) reaches its minimum value, -c:

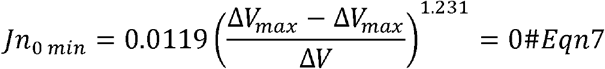

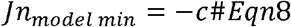

At this point, the corrected sap flux density (Jn_new_, cm s^-1^), calculated with the correct baseline (Baseline_new_) also reaches this minimum value -c:

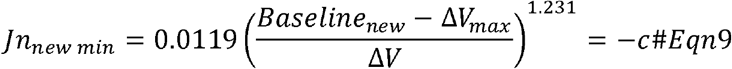

Solving for Baseline_new_ gives

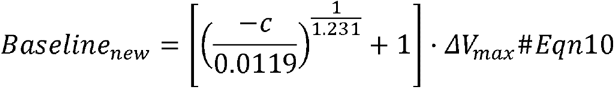

With Baseline_new_, the Jn_new_ can be calculated as:

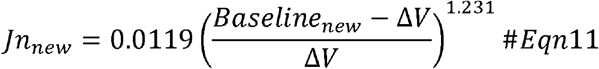

To ascertain whether transpiration or refilling could independently explain nighttime sap flux density, univariate regression models using only VPD or R_Δd_ were also constructed on a night-by-night basis for comparison:

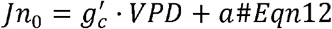

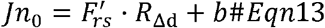

where g_c_’, F_rs_’, a, and b were constants within each night.

The suffixes n and d are used to partition between nighttime and daytime sap flux density (Jn and Jd), respectively. The subscript _0_ implies the sap flux density calculated with ΔV_max_ as the baseline, including Jn_0_ and Jd_0_. The subscript _model_ implies the sap flux density calculated based on transpiration and refilling (Eqn5), including Jn_model_. The subscript _new_ implies sap flux density related variables calculated with the new baseline derived from model (Eqn10), including Baseline_new_, Jn_new_ and Jd_new_. Refer to Table 1 for the full terminology used in the study.

**Table 1:** Summary of the terminology and definitions.

| Terminology | Units | Definition |
| --- | --- | --- |
| $T_{\text{measure}}$ | $\text{mg s}^{-1} \text{ m}^{-2}$ | Leaf transpiration rate. |
| ET | $\text{mm 30min}^{-1}$ | Evapotranspiration. |
| $R_{\Delta d}$ | $\mu\text{m s}^{-1}$ | Rate of change in stem diameter, indicating the amount of stem diameter change per unit of time. |
| $g_c$ | $\text{cm s}^{-1} \text{ kPa}^{-1}$ | Canopy conductance, modelled each night, represents the amount of sap flux used for transpiration driven by per unit of VPD. |
| $F_{rs}$ | Unitless | Refilling swelling factor, modeled each night, represents the amount of sap flux used for refilling caused by per unit of $R_{\Delta d}$. |
| $g_c'$ | $\text{cm s}^{-1} \text{ kPa}^{-1}$ | Assuming that nighttime sap flux is driven entirely by transpiration, and $g_c'$ represents the sap flux driven by per unit of VPD. |
| $F_{rs}'$ | Unitless | Assuming that nighttime sap flux is used entirely by refilling, and $F_{rs}'$ represents the sap flux that caused by per unit of $R_{\Delta d}$. |
| $\Delta V$ | mV | Voltage difference between the heated probe and reference probe of TDP. |
| $\Delta V_{\text{max}}$ | mV | Maximum value of $\Delta V$ for each night, serves as baseline to calculate $J_{n0}$. |
| Baseline<sub>new</sub> | mV | Baseline used to calculate $J_{n\text{new}}$, obtained from modelling (Methods S1). |
| $\Delta\text{Baseline}$ | % | Relative difference between Baseline<sub>new</sub> and $\Delta V_{\text{max}}$, i.e., $(\text{Baseline}_{\text{new}} - \Delta V_{\text{max}}) / \Delta V_{\text{max}} * 100\%$. |
| $J_{n0}$ | $\text{cm s}^{-1}$ | Nighttime sap flux density calculated based on $\Delta V$ and preliminary baseline ($\Delta V_{\text{max}}$ of each night). |
| $J_{n\text{new}}$ | $\text{cm s}^{-1}$ | Nighttime sap flux density calculated based on $\Delta V$ and true baseline (Baseline<sub>new</sub> of each night). |
| $J_{n\text{model}}$ | $\text{cm s}^{-1}$ | Nighttime sap flux density calculated based VPD and $R_{\Delta d}$ as well as the $g_c$ and $F_{rs}$ obtained from the model. |
| $J_{d0}$ | $\text{cm s}^{-1}$ | Daytime sap flux density calculated based on $\Delta V$ and preliminary baseline ($\Delta V_{\text{max}}$ of each night). |
| $J_{d\text{new}}$ | $\text{cm s}^{-1}$ | Daytime sap flux density calculated based on $\Delta V$ and true baseline (Baseline<sub>new</sub> of each night). |
| J | $\text{cm s}^{-1}$ | Daily mean sap flux density. |

The independent partitioning evaluation was conducted using linear regression models. Compared to Jn_0_, Jn_model_, and Jn_new_, T was expected to show strongest correlation with T_measure_ and ET, as it represents the portion of sap flux used only for transpiration, excluding water that remained in the stem for refilling. The homogeneity of the even aged Scots pine stand in Hyytiälä enables us to qualitatively extrapolate from individual trees to the ecosystem level (Vesala et al., 1998). Additionally, we conducted a residual analysis of the evaluation model, using root mean square error (RMSE) and mean absolute error (MAE) to assess residual deviation. It is important to note that the correlation analysis here were not designed for precise quantitative predictions of T_measure_ or ET but rather to qualitatively evaluate the partitioning, given the inherent noise in nighttime T_measure_ and ET data. Fig. 1 provides an overview of the measurements and theoretical framework.

**Fig. 1:**
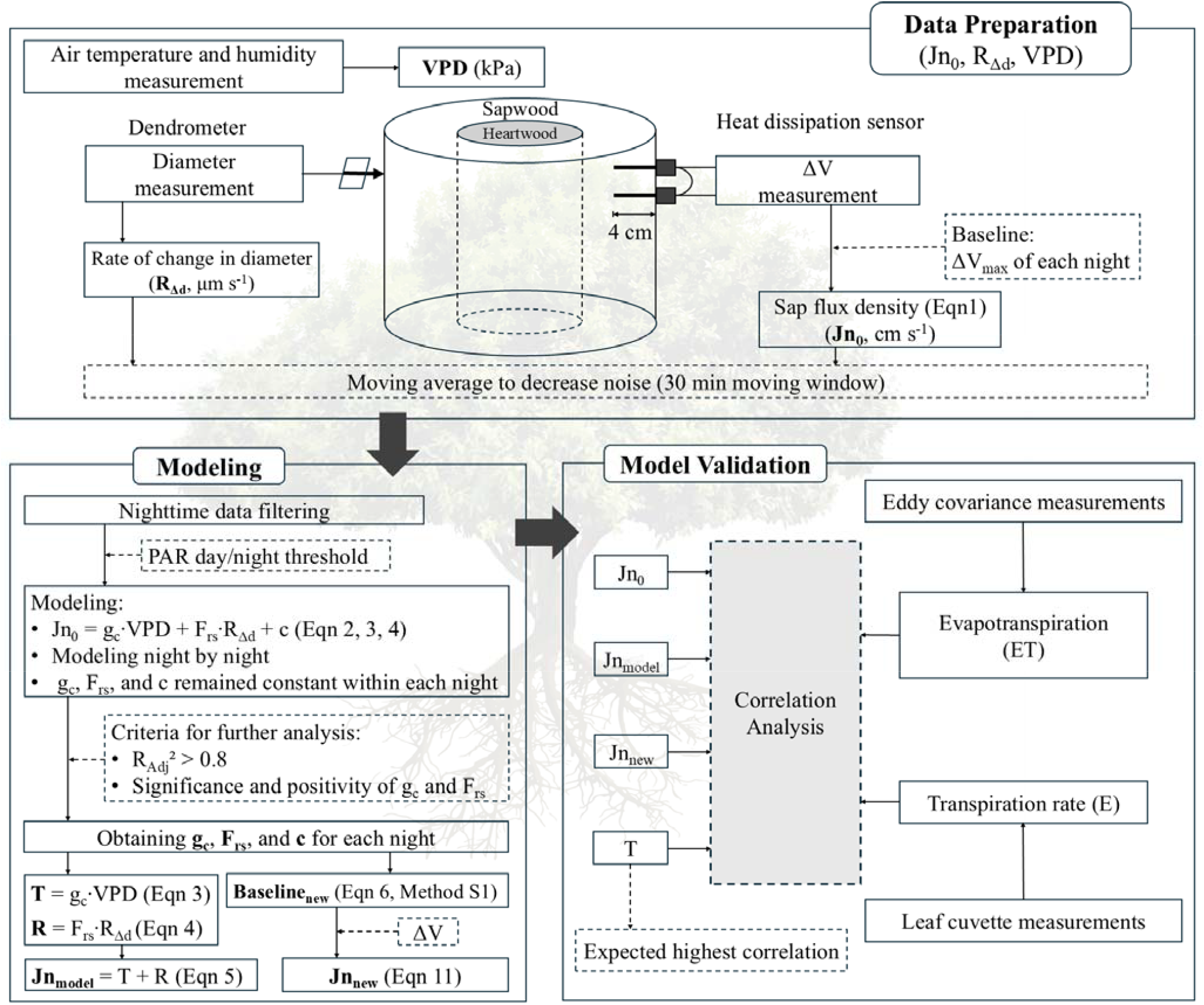
Measurements and theory. VPD represents vapor pressure deficit (kPa), R_Δd_ is the rate of change in diameter from dendrometer measurements (μm s□^1^), g_c_ denotes canopy conductance (cm s□^1^ kPa□^1^), and F_rs_ indicates the refilling swelling factor (unitless). Jn_0_ refers to nighttime sap flux density using ΔV_max_ as baseline (Eqn1), Jn_new_ represents sap flux density using Baseline_new_ as baseline (Eqn11), and Jn_model_ is the sap flux density obtained by the sum of transpiration and refilling flux density from the model (Eqn5).

### Statistical analyses

Both sap flux density and stem diameter data were resampled to a 5-minute resolution and then smoothed using a 30-minute moving average window. This process was undertaken to minimize noise while ensuring adequate number of data points were available each night for model fitting.

The Shapiro-Wilk test was conducted to examine the normality of the data. As the data was not normally distributed, the Kruskal-Wallis test was used to assess the significance of differences in R^2^ among different models and Jn values derived from different methods. Levene’s test was conducted to examine for homogeneity of variances of R_Adj_^2^. Pairwise comparisons were then performed using the Mann-Whitney U test. The t-test was employed to compare ΔV_max_ and Baseline_new_, as well as Jd and Jn/Jd based on different baseline. Pearson’s correlation was introduced to explore the relationship between model parameters. Akaike Information Criterion (AIC) was employed to assess different models. Loess fit was used to reveal the seasonality of VPD, T, R, g_c_ and F_rs_. Pandas and statsmodels packages in Python 3.11.5 were used for data preprocessing and nightly modelling, RStudio 2023.12.1 for statistical analysis and data visualization. Origin 2024 was also used for data visualization.

## Results

### Baseline problem

The intercept c was systematically negative in the multiple linear regression model (Eqn2) (Fig. S4), indicating that the actual baseline (Baseline_new_) was systematically higher than ΔV_max_ for both species (Fig. 2A, Table 2, Fig. S5A-C) (*P* < 0.01), by an average of 6.52% and 6.20% for *P. sylvestris* and *A. glutinosa*, respectively. Jn_model_ and Jn_new_ were largely overlapped in all years (Fig. 2B, Fig. S5D-F), and there was no significant difference between them (Table 2). However, both Jn_model_ and Jn_new_ were significantly higher than Jn_0_ (*P* < 0.01) (Table 2). Over the season, the average relative difference between nighttime Jn_new_ and Jn_0_ was 204.56% for *P. sylvestris* and 344.77% for *A. glutinosa* (Fig. 2B, Fig. S5D-F). Likewise, daytime sap flux density (Jd_0_) and ratio of Jn_0_ to Jd_0_ (Jn_0_/Jd_0_) were significantly underestimated when ΔV_max_ was used as the baseline (Eqn1) compared to those calculated with Baseline_new_ (Eqn11) (Jd_new_, Jn_new_/Jd_new_) (*P* < 0.01) (Fig. 2C and D, Table 2). This resulted in daytime Jd_new_ averaging 28.30% higher than Jd_0_ for *P. sylvestris* and 45.68% higher for *A. glutinosa* (Fig. 2C, Fig. S5G-I). Additionally, the Jn_0_/Jd_0_ were underestimated 132.22% (*P. sylvestris*) and 202.10% (*A. glutinosa*) compared to Jn_new_/Jd_new_ (Fig. 2D, Fig. S5J-L). The sap flux density calculated for the full-day (from sunrise to the next sunrise) using Baseline_new_ as a baseline was significantly higher than that calculated using ΔV_max_ as a baseline (*P* < 0.01), by an average of 45.20% for *P. sylvestris* and 54.76% for *A. glutinosa*.

**Table 2:** Comparison of baseline (Baseline, mV), daytime sap flux density (Jd, 10^-3^ cm s^-1^), nighttime sap flux density (Jn, 10^-3^ cm s^-1^), ratio of Jn to Jd (Jn/Jd), and full-day sap flux density (J, 10^-3^ cm s^-1^) across different methods of calculating sap flux density. Data shown: mean ± SD.

|  | *P. sylvestris* |  |  |  |  | *A. glutinosa* |  |  |  |  |
| --- | --- | --- | --- | --- | --- | --- | --- | --- | --- | --- |
|  | Baseline | Jd | Jn | Jn/Jd | J | Baseline | Jd | Jn | Jn/Jd | J |
| Baseline<sub>new</sub> | 0.25 ± 0.01 a | 2.40 ± 1.04 a | 0.73 ± 0.39 a | 0.31 ± 0.12 a | 1.91 ± 0.88 a | 0.70 ± 0.02 a | 2.37 ± 0.70 a | 0.58 ± 0.20 a | 0.25 ± 0.08 a | 1.94 ± 0.59 a |
| $\Delta V_{\max}$ | 0.23 ± 0.01 b | 1.90 ± 0.78 b | 0.25 ± 0.10 b | 0.15 ± 0.09 b | 1.42 ± 0.60 b | 0.66 ± 0.02 b | 1.66 ± 0.50 b | 0.15 ± 0.07 b | 0.10 ± 0.05 b | 1.29 ± 0.42 b |
| model-based | - | - | 0.68 ± 0.35 a | - | - | - | - | 0.54 ± 0.18 a | - | - |

Baseline<sub>new</sub> indicates method of calculating sap flux density using Baseline<sub>new</sub> as baseline. $\Delta V_{\max}$ indicates method of calculating sap flux density using $\Delta V_{\max}$ as baseline. Model-based indicates sap flux density derived from model (Eqn5). Different lower-case letters indicate significant differences between methods ( $P < 0.01$ ).

**Fig. 2:**
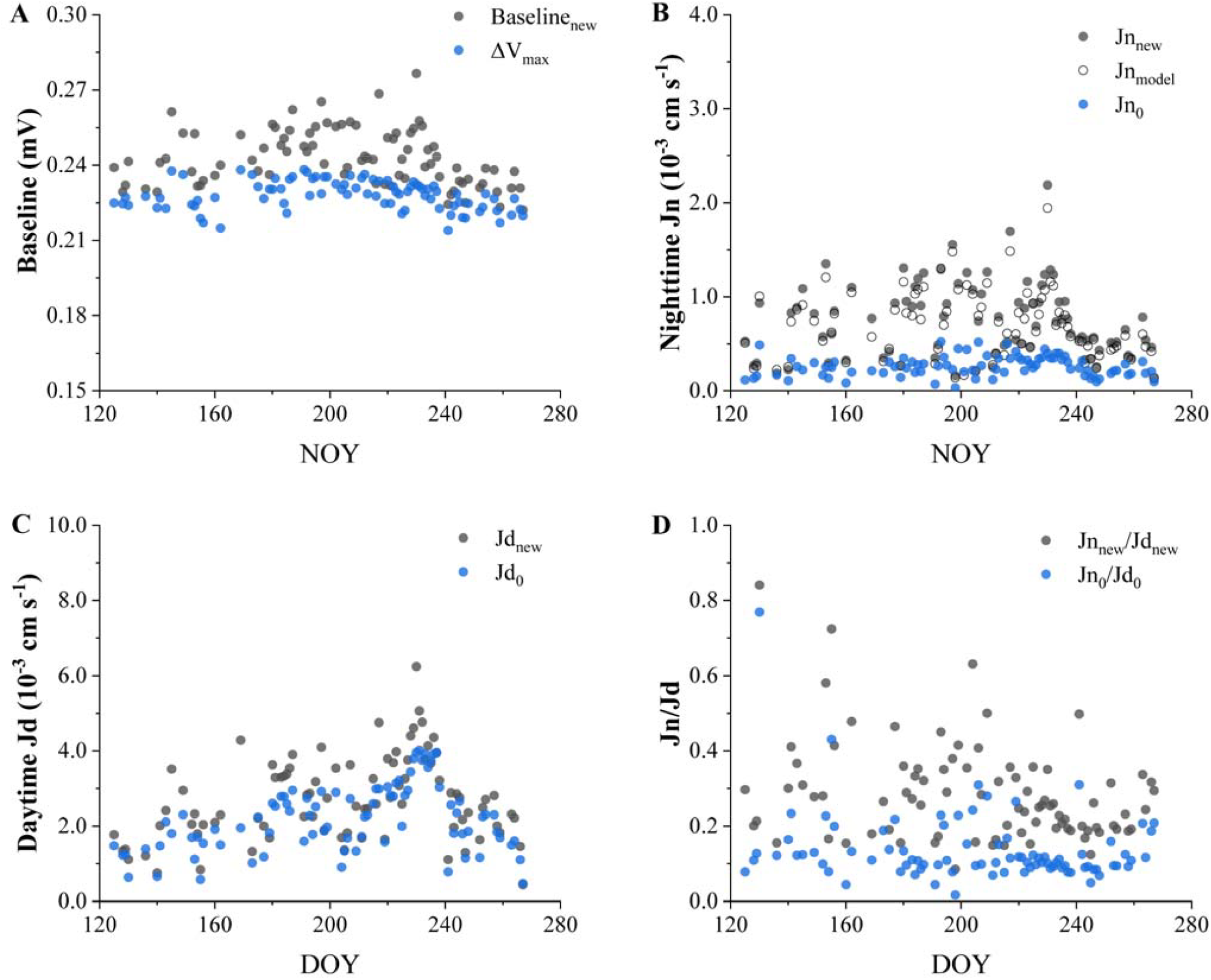
Baseline and related sap flux density of *P. sylvestris* in 2015. A) ΔV_max_ and model-obtained baseline (Baseline_new_) (Eqn10). B) Nighttime sap flux density with different baseline (Jn_0_, Jn_new_) (Eqn1, Eqn11), as well as the sap flux density derived from model (Jn_model_) (Eqn5). C) daytime sap flux density with different baseline (Jd_0_, Jd_new_) (Eqn1, Eqn11). D) Ratio of nighttime sap flux density to daytime with different baseline (Jn_0_/Jd_0_, Jn_new_/Jd_new_). Data resolution: 1 night or day.

At the lowest sap flux density each night (most likely to occur at the end of the night), VPD and R_Δd_ explained over 80% of the variation in difference between ΔV_max_ and baseline_new_ (ΔBaseline) for both tree species (Table 3). The increase in VPD or R_Δd_ led to an increase in ΔBaseline. This indicates that the higher the sap flux density attributed to transpiration and refilling, the larger the error caused by the original baseline. Alternatively, without using dendrometer, VPD, g_c_’, or the nightly mean Jn_0_ were used to build the model with ΔBaseline, and the intercept was also introduced to indirectly represent refilling. This explained over 50% of the ΔBaseline variation for both species and can thus be used to calibrate the baseline without R_Δd_ to some extent.

**Table 3:**
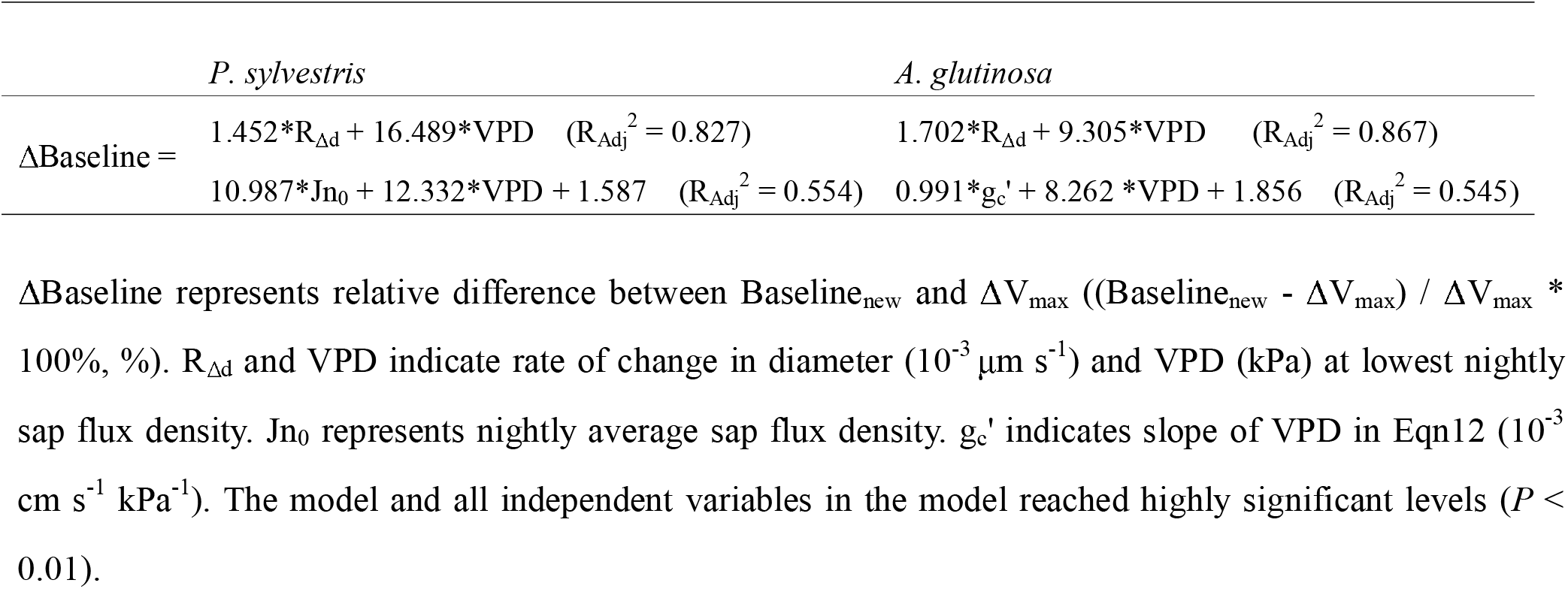
Modelling relative difference between Baseline_new_ and ΔV_max_ in relation to other variables for calibration.

There were differences in the relationship between R_Δd_ of *P. sylvestris* and Jn obtained by different methods (Fig. 3A-C). R_Δd_ had positive value when Jn_0_ was 0 (Fig. 3A), indicating that the sap flux density was underestimated due to stem refilling. This was not the case for Jn_new_ and Jn_model_ (Fig. 3B, C). Although Jn_new_ and Jn_model_ showed systematic improvement compared to Jn_0_, their patterns relative to R_Δd_ remained largely unchanged. Moreover, R_Δd_ could not independently explain Jn, i.e., their relationship was not entirely linear (Fig. 3A-C). In other years, *P. sylvestris* and *A. glutinosa* showed similar results (Fig. S6).

**Fig. 3:**
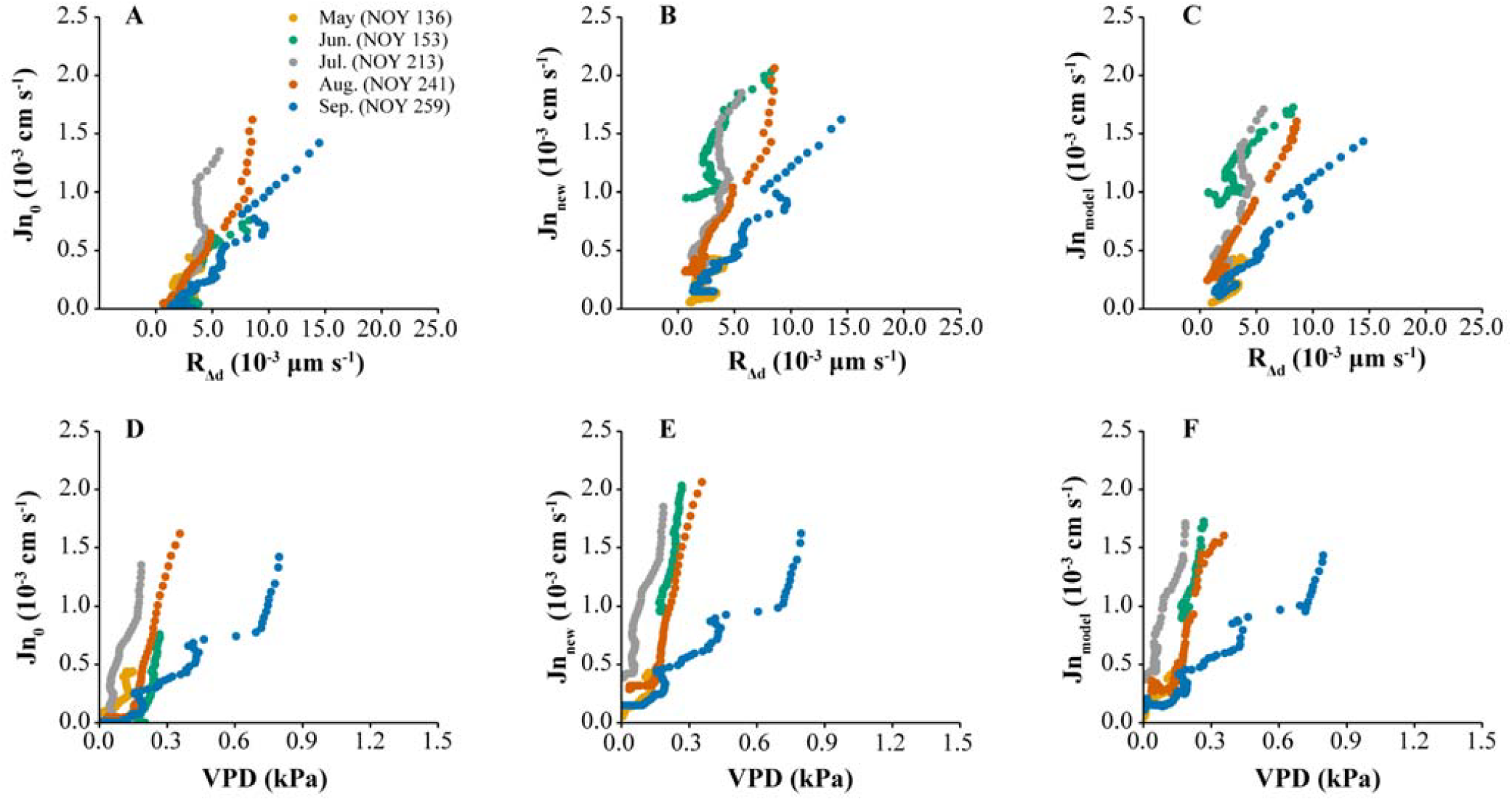
Rate of change in diameter (R_Δd_) of *P. sylvestris* and VPD in relation to sap flux density calculated using different ways in five individual nights in 2015. Jn_0_ and Jn_new_ indicate nighttime sap flux density using ΔV_max_ as baseline (Jn_0_) (Eqn1) and using Baseline_new_ as baseline (Jn_new_) (Eqn11), and sap flux density derived from model (Jn_model_) (Eqn5). Data resolution: 5 min.

Similar results were found regarding the relationship between VPD and Jn (Fig. 3D-F, Fig. S7). Fig. S8 and S9 offer entire data for both tree species in each year, mirroring the results obtained from five individual nights (Fig. 3).

### Partitioning between transpiration and refilling

On most nights, the model (Eqn2) successfully partitioned between transpiration and refilling (on average 70.83% of total nights) (Table S1). However, in some cases the reliability of the model parameters was not meaningful and therefore model on these nights was not used for calculating T, R, and Baseline_new_ (on average 29.17% of total nights) (Table S1, Fig. S10). Specifically, rainfall affected measurement, making 13.96% of the nights too noisy. R_Δd_ and VPD had a highly similar trend on 4.17% of nights, which caused covariance problems in the model (*P* > 0.05). 3.96% of the nights were not fitted due to noise of dendrometer, potentially caused by errors. Additionally, model did not work well on only 0.63% of nights because either transpiration or refilling was too minimal. Finally, on 6.46% of nights the data was not modelled properly due to an unspecified reason.

### Evaluation of partitioning

The R_Adj_^2^ of the univariate regression models using only VPD or R_Δd_ (Eqn12 or Eqn13) were significantly lower than that of the multiple linear regression model incorporating both (Eqn2) (Table S2). The variability of the R_Adj_^2^ of the univariate regression models (Eqn12 or Eqn13) was significantly greater than that of the multiple linear regression model (Eqn2). In contrast, the multiple linear regression model (Eqn2) consistently maintained a high R_Adj_^2^ across most nights with lower AIC (Table S2, Fig. S11).

Jn_0_, Jn_new_, Jn_model_, and T were significantly positively correlated with nighttime T_measure_ (*P* < 0.05), with T having the highest correlation (Fig. 4). Similarly, Jn_0_, Jn_new_, and Jn_model_ did not correlate with nighttime ET (*P* > 0.05) (Fig. 5A-C), but T and nighttime ET were significantly positively correlated (*P* < 0.01) (Fig. 5D). Furthermore, T had the lowest RMSE and MAE among the models (Figs. 4H and 5H), indicating smaller residual deviations. These results were not meant for precise quantitative prediction of ET or T_measure_, but rather to qualitatively evaluate the effectiveness of the partitioning method. The better performance of T indicates that it successfully represents the portion of sap flux used for transpiration, excluding the portion used for refilling (R). Additionally, many instances were observed where T_measure_ and ET was high while Jn_0_ was close to 0 (Fig. 4A, 5A), indicating that the original baseline (ΔV_max_) and sap flux density based on it (Jn_0_) were underestimated.

**Fig. 4:**
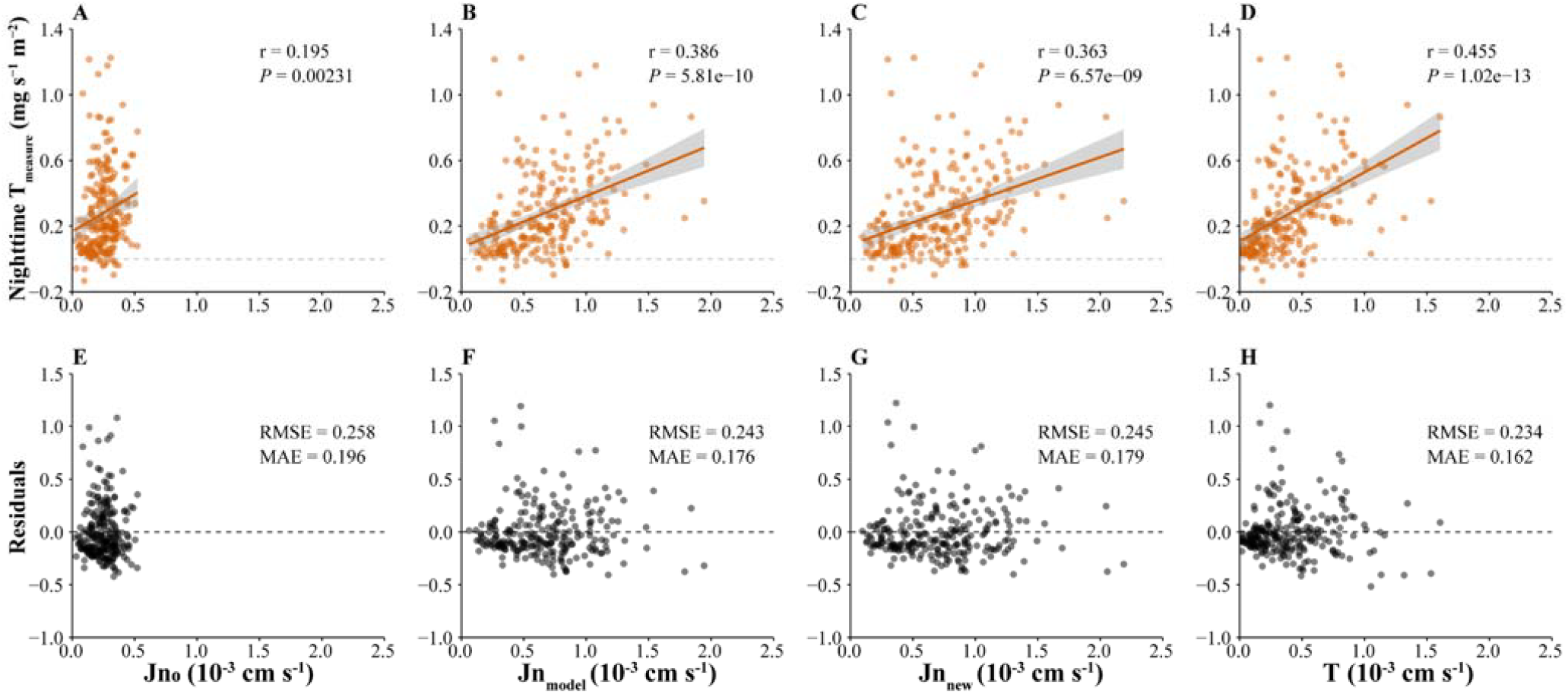
Relationship between nighttime transpiration rate (E) and sap flux density. A) Nighttime sap flux density calculated using ΔV_max_ as baseline (Jn_0_), B) nighttime sap flux density calculated from model (Jn_model_), C) nighttime sap flux density calculated using Baseline_new_ as baseline (Jn_new_), and D) model-calculated nighttime sap flux density used for transpiration (T) in *P. sylvestris*. E-H) Corresponding residuals. Data resolution: 1 night.

**Fig. 5:**
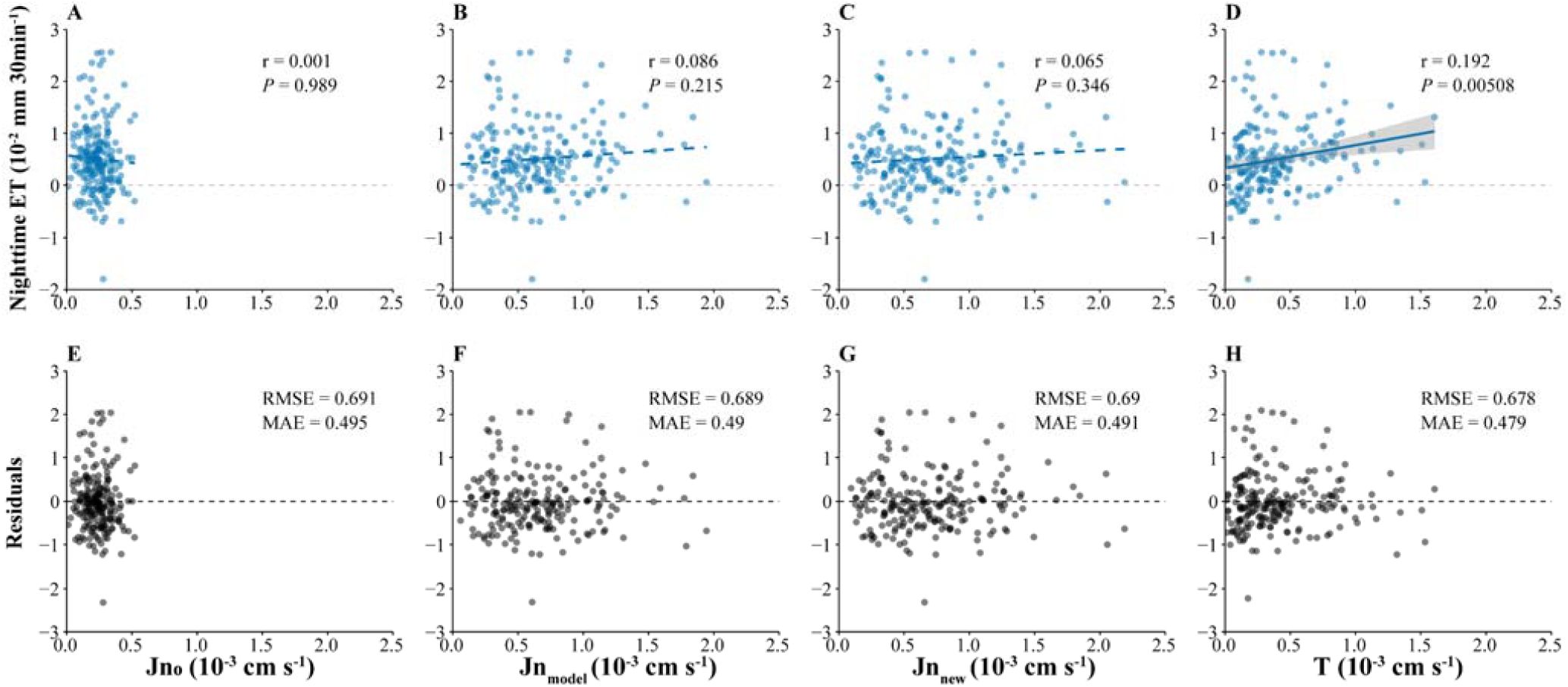
Relationship between nighttime ET and sap flux density. A) Nighttime sap flux density calculated using ΔV_max_ as baseline (Jn_0_), B) Nighttime sap flux density calculated from model (Jn_model_), C) Nighttime sap flux density calculated using Baseline_new_ as baseline (Jn_new_), and D) model-calculated nighttime sap flux density used for transpiration (T) in *P. sylvestris*. E-H) Corresponding residuals. Data resolution: 1 night.

Jn_0_, R_Δd_, and VPD typically decreased during nights (Fig. 6). Jn_0_ decreased gradually, and Jn_model_ consistently tracked the trend of Jn_0_ but systematically exceeded it. R_Δd_ and VPD showed fluctuating decreases, with their trends mirroring each other (Fig. 6). This suggests a multiple linear regression relationship among them, with relatively fixed constants (g_c_ and F_rs_) each night. Interestingly, there was almost never such a moment during the night when both VPD and R_Δd_ reached 0 (Fig. 6).

**Fig. 6:**
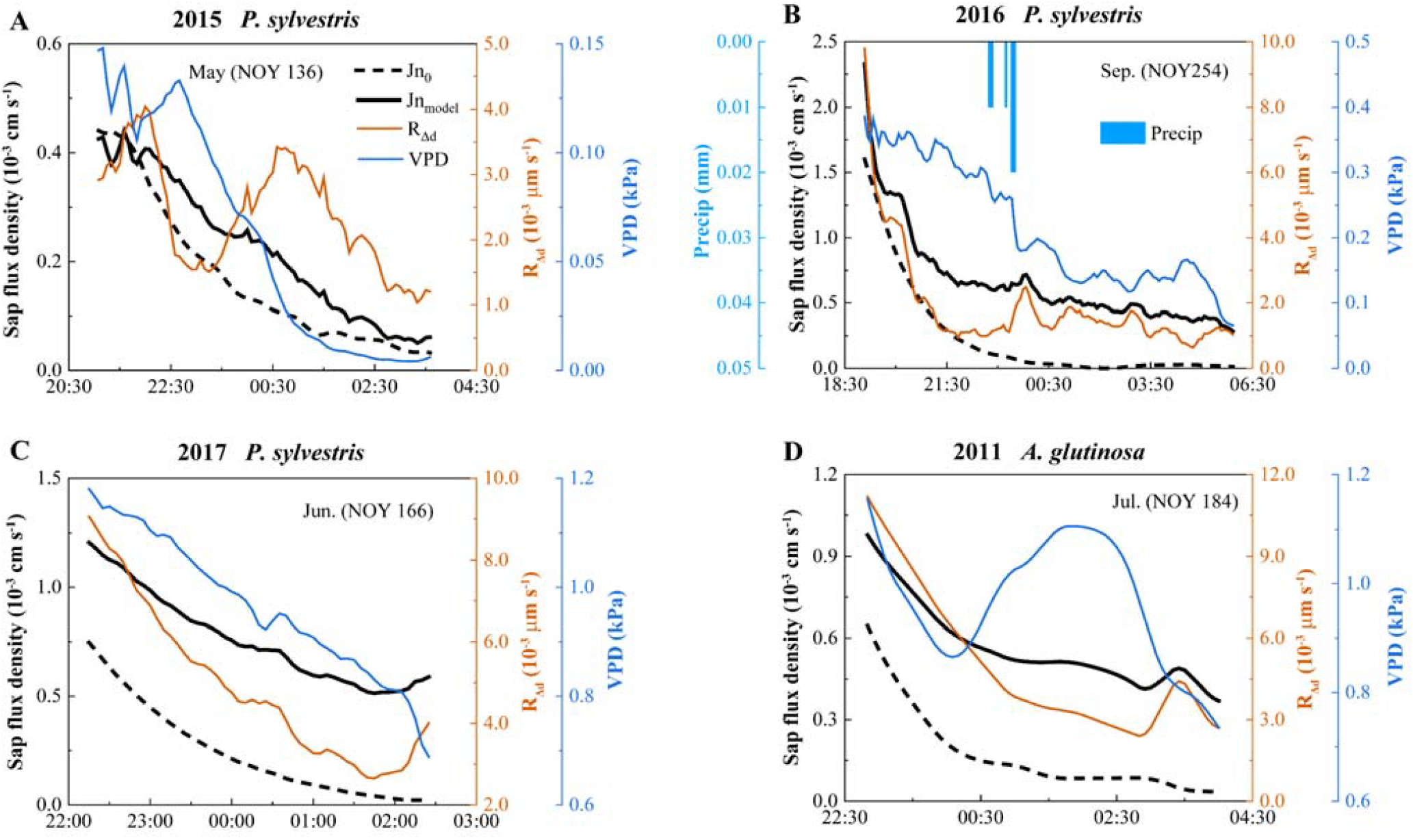
Nighttime temporal trend in sap flux density, rate of change in diameter (R_Δd_), VPD, and precipitation (Precip). Jn_0_ and Jn_model_ indicate sap flux density calculated using ΔV_max_ as baseline (Eqn1) and that calculated from model Jn_model_ (Eqn5), respectively. Data resolution: 5 min.

There was no correlation between g_c_ and F_rs_ as the coefficients of VPD and R_Δd_ in the multiple linear regression model (Eqn2), while they were significantly correlated with g_c_’ and F_rs_’ in the univariate regression model (Eqn12 or Eqn13) (Table 4). It is noteworthy that g_c_’ was not only positively correlated with g_c_ (*P* < 0.01), but also positively correlated with F_rs_ (*P* < 0.01) (Table 4). Similarly, a significant positive correlation of F_rs_’ with g_c_ was also found (*P* < 0.01) (Table 4). This indicates that Jn explained by the univariate model of either VPD or R_Δd_ (Eqn12 or Eqn13) integrates the effects of another process.

**Table 4:**
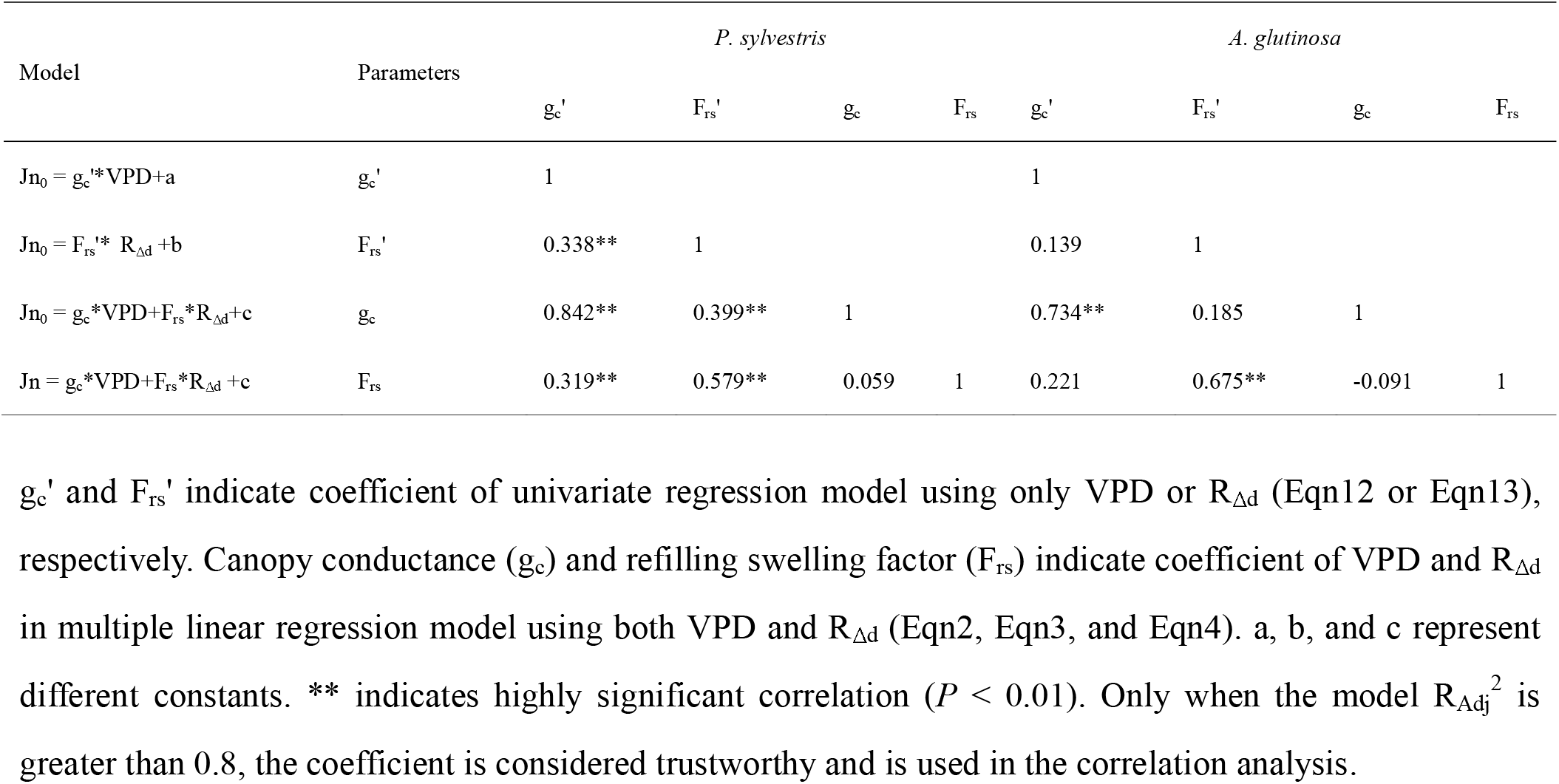
Correlation of coefficient for different models fitting nighttime sap flux density.

### Transpiration and refilling pattern

The ratios of T and R to total nightly sap flux density (Jn_model_) show their seasonal characteristics (Fig. 7). For *P. sylvestris*, R/Jn_model_ progressively surpassed T/Jn_model_ after NOY 200, indicating a shift where refilling became dominant over transpiration, primarily in August (Fig. 7A-C). In contrast, *A. glutinosa* showed stable T/Jnmodel and R/Jnmodel ratios throughout the observation period, with no similar seasonal shift (Fig. 7D). Additionally, T/Jnmodel for *P. sylvestris* began to decline after a peak in June, whereas *A. glutinosa* did not exhibit this decline by late July, when observations ended (Fig. 7D). Fig. S12 also shows the time series of the absolute values of T and R.

**Fig. 7:**
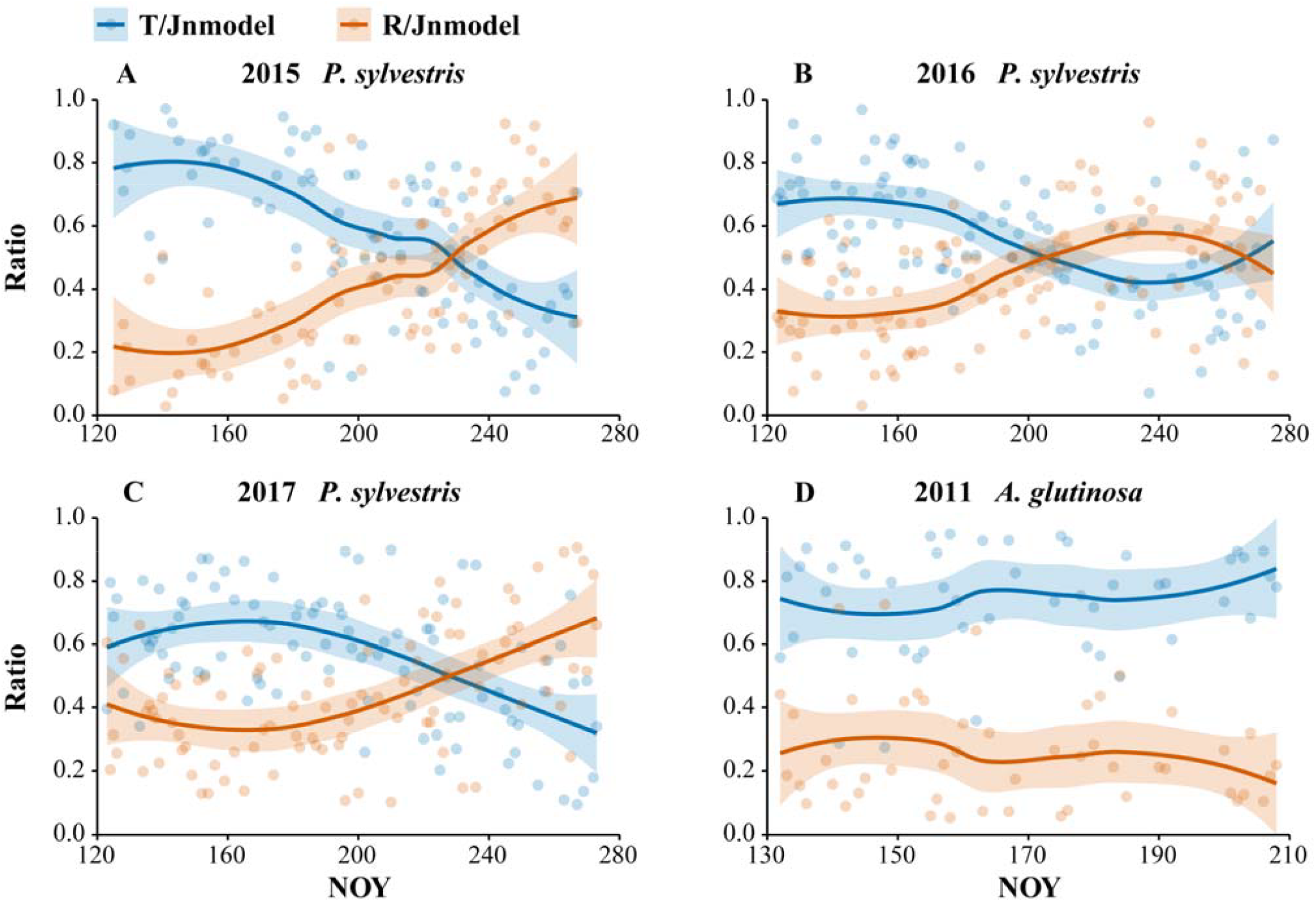
Time-series in ratio of transpiration flux (T) and refilling flux (R) to total nighttime sap flux density (Jn_model_).

## Discussion

### Partitioning between transpiration and refilling

The time-separation approach and its derivatives have been proposed to partition between refilling and transpiration (Fisher et al., 2007; Liu et al., 2020). These methods sometimes result in scenarios where only refilling or only transpiration is observed for a period of time. In our study, however, on almost all nights, either VPD or R_Δd_ failed to align the trend of Jn and independently explain it well enough (Fig. 3, 6, S8, S9). Conversely, the multiple linear regression model (Eqn2) combining both VPD and R_Δd_ explained scenarios that the univariate models (Eqn12 or Eqn13) couldn’t capture, thereby improving R_Adj_^2^ and reducing its variability (Table S2, Fig. S11). Since the multiple regression model statistically ensures that both VPD and R_Δd_ contribute simultaneously, these results suggest that refilling and transpiration almost always occurred concurrently throughout the night, rather than sequentially. Similarly, efforts to partition refilling and transpiration through time series modelling have relied solely on the goodness of fit to determine the success of the distinction (Zhao et al., 2022). However, a priori modelling may be more convincing only after empirical evaluation. For example, our findings indicate that univariate models utilizing only VPD or R_Δd_ Eqn12 or Eqn13) can still account for a considerable variation of Jn in certain instances, with their overall R_Adj_ ^2^ typically exceeding 0.6 (Table S2, Fig. S11). This is because on some nights, either refilling or transpiration dominated to the extent that ignoring the other could be statistically acceptable. Additionally, the collinearity between R_Δd_ and VPD means that either one can integrate the other in the model, resulting in a high R_Adj_ ^2^, even if obtained results are biased. This is confirmed by the significant positive correlation found between F_rs_’ and g_c_, as well as between g_c_’ and F_rs_ (Table 4). Therefore, F_rs_’ and g_c_’ were overestimated in the univariate regression models (Eqn12 and Eqn13) compared to F_rs_ and g_c_ in the multiple linear regression model (Eqn2), and the direct relationship between VPD and Jn could be biased.

Based on real measurements, some efforts have aimed to use leaf gas exchange measurements to obtain true transpiration, where the difference between total sap flow and transpiration represents the refilling component (Wang et al., 2012). While transpiration can be obtained through leaf gas exchange measurements, a challenge lies in the assumption that individual leaves represent the transpiration of the entire canopy. Additionally, gas exchange measurements during the nighttime are technically difficult to conduct when RH is high due to condensation of water in the chambers and inlets and outlets (Brito et al., 2018; Dawson et al., 2007; Escalona et al., 2013). Calculating refilling based on differences in sap flow at different tree heights offers additional insights (Zeppel et al., 2010). Although thermal diffusion probes effectively capture relative amounts and dynamic trends in sap flux density, converting these measurements to absolute values can introduce systematic errors. These errors can arise e.g. from spatial and temporal variations in wood thermal and hydraulic properties (Hölttä et al., 2015; López-Bernal et al., 2014), differences in microenvironments, and even variations between individual sensors (Oishi et al., 2016).

No solution is perfect, and the model did not perform well on 29.17% of the total nights in our study (Table S1, Fig. S10). In our study, rainfall was the primary environmental factor negatively impacting measurements, with nearly half of the model failures attributed to this issue (Table S1). Rain posed an inevitable challenge for field measurements, particularly for stem diameter measurements, as it causes stems to swell by absorbing rainwater and then shrink as the rainwater evaporates. Since these changes in diameter due to rainwater are not contributed by the sap flux density, this poses a problem for modelling. Additionally, the collinearity between R_Δd_ and VPD presented a statistical disadvantage in the modelling, as both variables exhibited similar temporal patterns throughout each night (Table S1). After sunset, VPD declines as air temperature drops and relative humidity (RH) rises, while stem swelling slows down due to a reduced water potential difference between the soil and the stem. Moreover, the TDP has known limitations in accurately measuring low sap flow rates (Flo et al., 2019). Depending on the species and environment, using alternative sensor types, such as heat pulse sensors, may help improve the model’s performance.

### Nighttime transpiration of trees induced evapotranspiration response

At the SMEAR II site, the response of nighttime ET induced by T was observed (Fig. 5), thereby validating the model that partitions between transpiration and refilling. Similarly, Fisher et al. (2007) attempted to explore the relationship between nighttime sap flow and ET: while there were instances during the year when they appeared to follow the similar trend, most of the time no correlation was found between nighttime sap flow and ET. The main reasons for the differences between our study and theirs are as follows: (1) the forests at the SMEAR II site in our study are equal-aged and homogeneous *P. sylvestris* (Vesala et al., 1998). Consequently, ET induced by tree transpiration was less prone to error due to heterogeneous individual trees and exhibited less dynamic variation. (2) Sap flow comprises both transpiration and refilling. Water refilling into the stems does not evaporate and therefore does not elicit response of nighttime ET, so the relationship between sap flow and ET can be biased. (3) Although sap flow at the oak-savanna site in the study by Fisher et al. (2007) exhibited a somewhat similar nightly pattern to ET, the seasonal increase of ET in summer was not reflected in sap flow, which was consistently near zero. As concluded by Fisher et al. (2007), nocturnal sap flow in their study could be underestimated.

### Nightly and seasonal dynamics of transpiration and refilling

Seasonally, both T and R for the two species increased during the summer, with *P. sylvestris* subsequently showing a downward trend (Fig. 7, S12). T showed a positive correlation with seasonal VPD changes (r = 0.58, *P* < 0.01 for *P. sylvestris*; r = 0.34 and *P* < 0.05 for *A. glutinosa*), with higher T during summer VPD peaks that gradually decreased into autumn (Fig. 7, S1, S12). R was linked to daytime sap flux density (Jd_new_) (r = 0.59, *P* < 0.01 for *P. sylvestris*; r = 0.29 and *P* < 0.05 for *A. glutinosa*) (Fig. 7, 2C, S5, S12), where greater summer depletion of stem water during the day led to increased R on subsequent nights. However, both species may respond differently to factors such as soil water availability, wood properties, and the turgor-dependent stem growth (Oliva Carrasco et al., 2015; Zhao et al., 2022), resulting in variations in the timing of their peaks (Fig. 7, S12).

Interestingly, successful model performance suggests that both canopy conductance (g_c_) and the refilling swelling factor (F_rs_) could be considered constant within each night, though they varied from night to night. On the one hand, g_c_ during the day fluctuated largely because of the stomatal response to light, whereas at night there was no such response; on the other hand, g_c_ during the night also involved conduction through the bark and cuticle, which remained relatively stable, allowing g_c_ to be treated as a constant. In addition, nightly variations in F_rs_ indicate that the same rate of refilling flux in different night may result in different degrees of stem swelling. This implies that stem might adjust their nightly refilling water allocation strategies—for instance, directing more water to the apoplastic xylem might cause less detectable stem expansion, making F_rs_ appear higher.

### Baseline strategies

There is a classical scheme of using the ΔT_max_ (ΔV_max_ in our case) per night as the baseline, i.e., assuming zero sap flow each night (Zhu et al., 2017), or its derivatives include a moving average of ΔT_max_ over multiple days as a baseline (Lu et al., 2004; Rabbel et al., 2016). However, those methods inevitably lead to underestimation of both baseline and Jn as ΔT_max_ does not guarantee zero-flow. Our results also show that baselining with daily ΔV_max_ causes a significant underestimation of both daytime and nighttime sap flux density (Table 2), highlighting the importance of an accurate baseline in correctly estimating nighttime flux contributions, which in turn influences our understanding of plant water use and storage dynamics. Alternatively, environmental dependent method for zero-flow condition is when evaporative demand (VPD) falls below a certain threshold and ΔT stabilizes (Chen et al., 2020; Oishi et al., 2016, 2008). Some efforts have been made to define “low evaporative demand” and “stable ΔT” more rigorously to increase the validity of baselining. For example, approaches have included considering both VPD and wind speed thresholds to determine a small transpiration drive (Zeppel et al., 2010), or using the ratio of nightly variation to daily variation in ΔT over a weekly moving window to ascertain whether “stability” has been achieved (Oishi et al., 2016). However, these stricter criteria make it more challenging to identify times when the conditions are met in practice, especially in boreal regions summer, where the nights are short, and there are so many nights when the criteria are not fulfilled. Another alternative method is to directly fit ΔT and VPD, using the intercept of the fitted curve when VPD equals to 0 as a baseline (Meinzer et al., 2013). While this approach neglects refilling, it could be enhanced by incorporating stem diameter parameters. Hence, the night-by-night use of R_Δd_ and VPD, which accounts for both refilling and transpiration, presents a promising strategy for enhancing the validity and reliability of long-term baselining.

The baseline error (ΔBaseline) between ΔV_max_ and the true baseline (Baseline_new_) largely originates from both transpiration and refilling processes and can thus be represented by VPD and R_Δd_ (Table 3). While incorporating dendrometer data allows refilling to be accounted for in baselining, the independent use of thermal diffusion probes to monitor sap flux density remains predominant. Therefore, we provide calibration models for *P. sylvestris* and *A. glutinosa* that do not rely on dendrometer data but use commonly available variables (Table 3). Jn_0_ and g_c_’ can partially represent R_Δd_ indirectly, as they both integrate refilling. It is worth noting that our method focuses on the case where zero-flow conditions are difficult to find or not present at all. Other calibrations such as dampening and sensor geometry calibration could also be considered to accurately quantify sap flow (Flo et al., 2019; Fuchs et al., 2017; Peters et al., 2018).

## Conclusion

Our study introduces an independently evaluated framework for partitioning between transpiration and refilling processes and proposes a practical baselining strategy based on this partitioning. The gradual dominance of R indicates a seasonal shift in nighttime sap flux allocation from transpiration to refilling, while variations in F_rs_ across nights suggest refilling water management strategies within stem tissues. As for TDP, potential use of baselining strategy based on VPD and dendrometers could be considered in calculating sap flux density for both daytime and nighttime across various environments and species. Moving forward, the partitioning between nighttime transpiration and refilling could be extended beyond TDP to other sensor types. Future work on the physiological and environmental drivers of nighttime transpiration and refilling will be important to advance our understanding of tree water use and ecosystem function in a hanging climate.

## Acknowledgements and Funding

We are thankful to Juuso Tuure for providing meteorological data in urban site and Anu Riikonen for providing the data from the *A. glutinosa* site. Support from the Academy of Finland Center of Excellence in Tree Biology (#342930) is acknowledged. MW acknowledges the support of China Scholarship Council program (NO. 202306510010). We appreciate Che Liu for the valuable comments on data analysis.

## Author contributions

TH conceptualized and designed the research. MW and MR performed the data analysis. MW wrote the manuscript with contributions from co-authors.

## Supplementary material

Additional supporting information may be found in the online version of this article.

### Methods S1 derivation process of Baseline_new_

**Fig. S1** Environmental background.

**Fig. S2** T-VPD time lag test.

**Fig. S3** Time series of nighttime canopy conductance (g_c_) and refilling swelling factor (F_rs_).

**Fig. S4** The intercept c of multiple linear regression model.

**Fig. S5** Baseline and related sap flux density of *P. sylvestris* and *A. glutinosa*.

**Fig. S6** Rate of change in diameter (R_Δd_) in relation to sap flux density derived from different methods in five individual nights.

**Fig. S7** VPD in relation to sap flux density derived from different methods in five individual nights.

**Fig. S8** Rate of change in diameter (R_Δd_) in relation to sap flux density derived from different methods.

**Fig. S10** Reasons for not being modelled well.

**Fig. S11** R_Adj_^2^ of different models.

**Fig. S12** Transpiration (T) and refilling (R) flux density.

**Table. S1** Performance distribution of model across nights

**Table. S2** Goodness of fit (R_Adj_^2^) of different models fitting nighttime sap flux density

