## Supplementary material for "Partitioning of nighttime transpiration and stem water refilling using VPD and dendrometer data: insights into baselining and nighttime sap flux interpretation"

**Author for correspondence:**

Mianzhi Wang

**Fig. S1** Soil volumetric water content (SWC), precipitation (Precip), as well as daytime and nighttime VPD in Hyytiälä from 2015 to 2017 and Helsinki in 2011. Data resolution: 1 night or day.

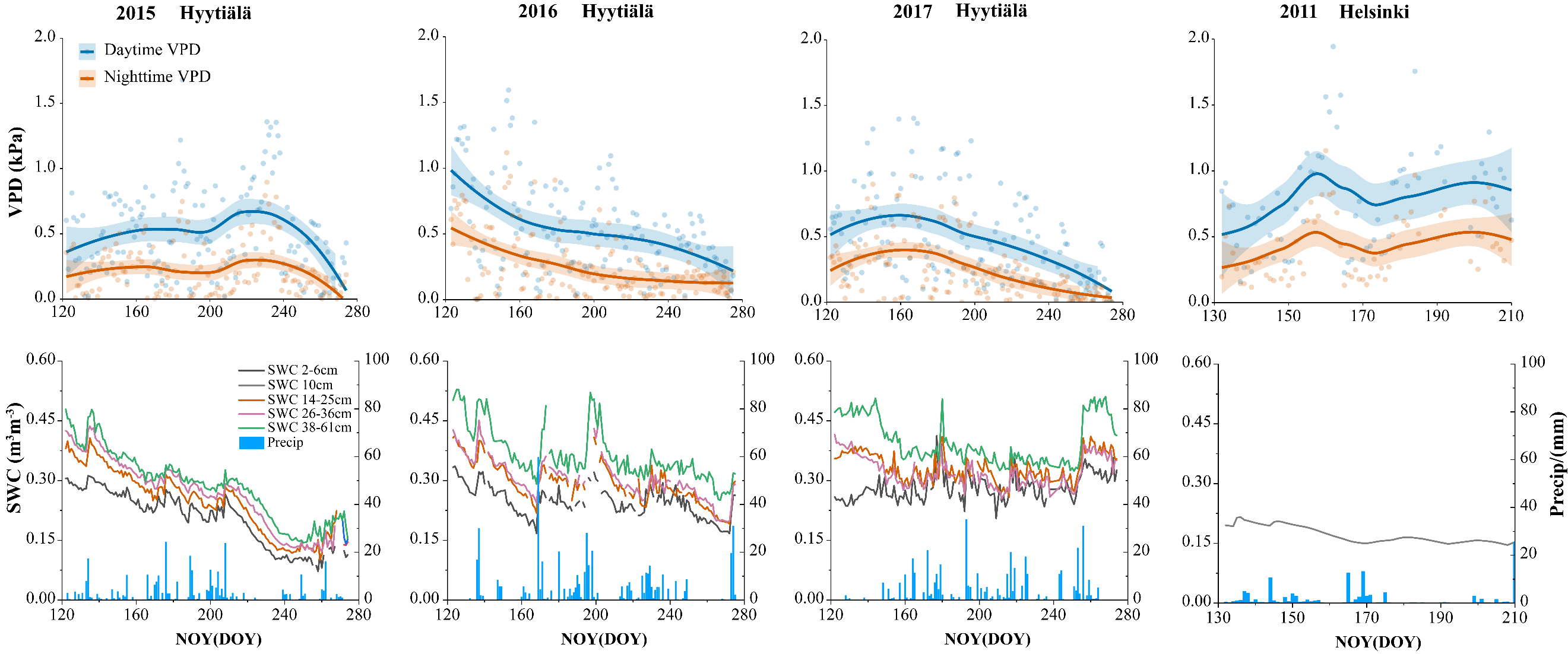

**Fig. S2** T-VPD time lag test. The timestamps of VPD were shifted backward by 5, 10, 15, 20, 30, 45, 60, 75, 90, and 120 minutes to account for a potential delay in T response to VPD. The model (Eqn2-4) was rerun to determine whether the shifted dataset provided a better fit. If a time lag were present, the adjusted R² (R_Adj_^2^, black) and Akaike Information Criterion (AIC, blue) values would indicate an optimal shift. The black and blue dashed lines represent the lowest R_Adj_^2^ and the highest AIC, respectively.

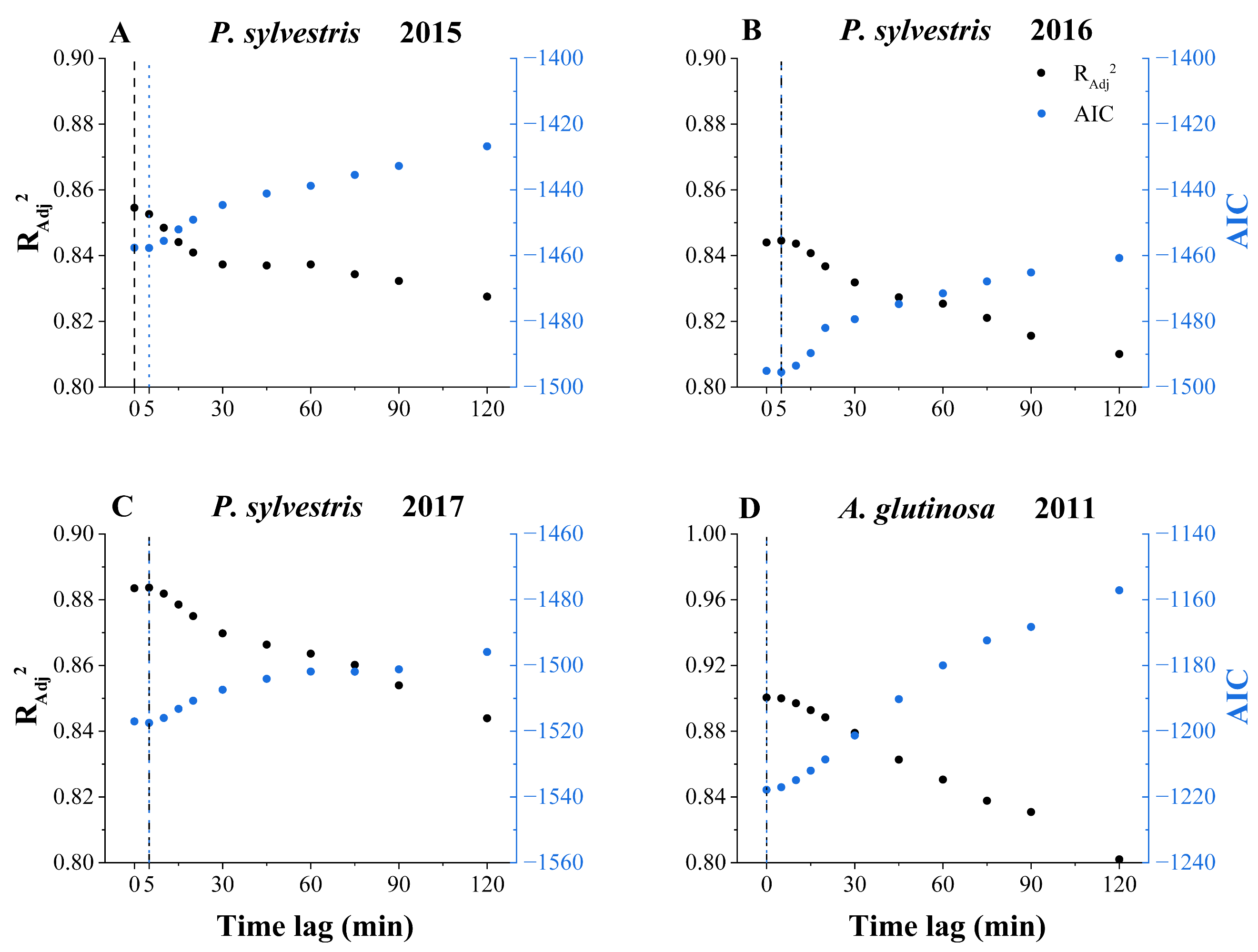

**Fig. S3** Time series of nighttime canopy conductance (g_c_) and refilling swelling factor (F_rs_). Data resolution: 1 night.

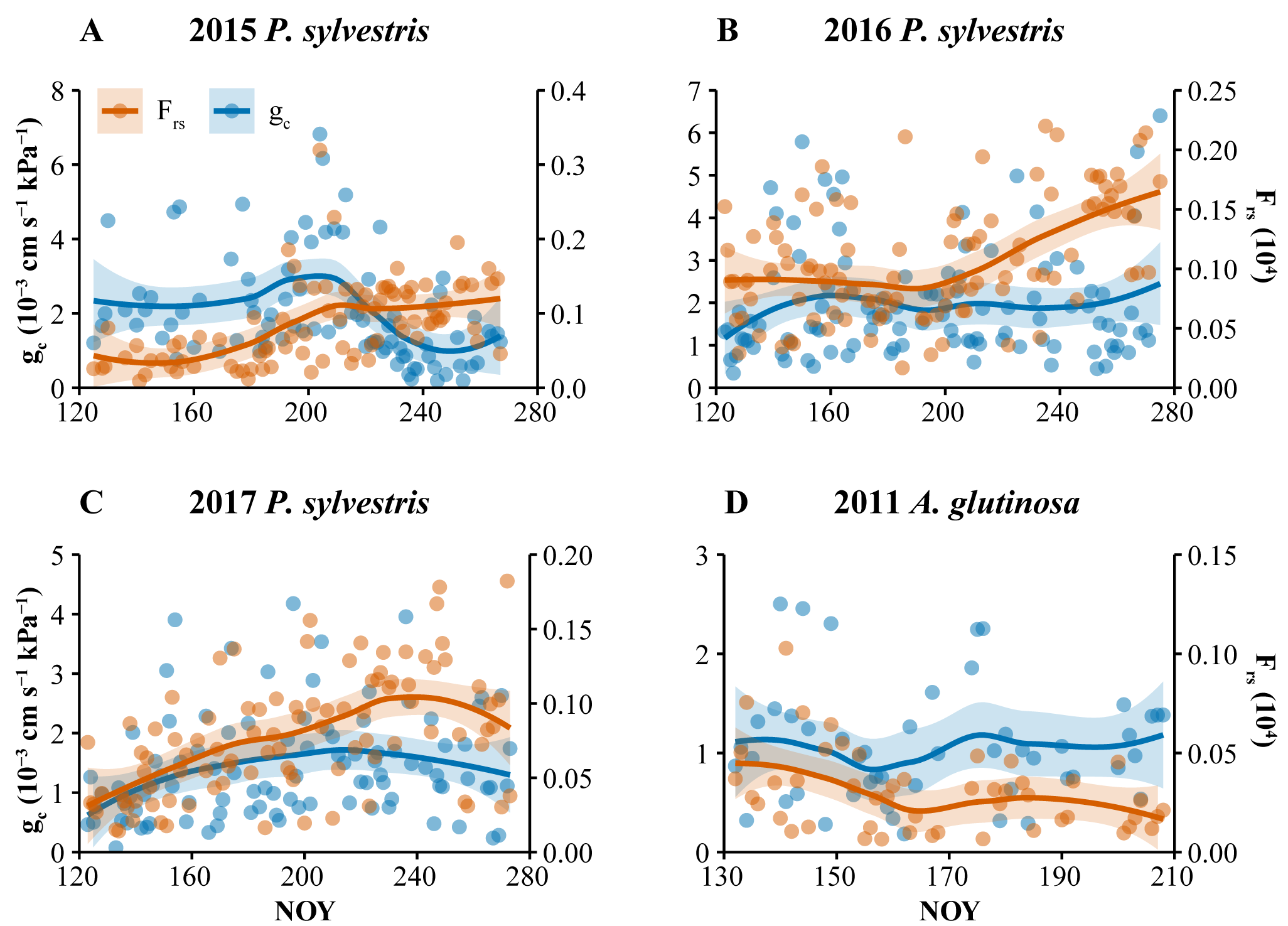

**Fig. S4** The intercept c of multiple linear regression model. The cases when c is greater than 0 was not used in the calculation of Baseline_new_. Data resolution: 1 night.

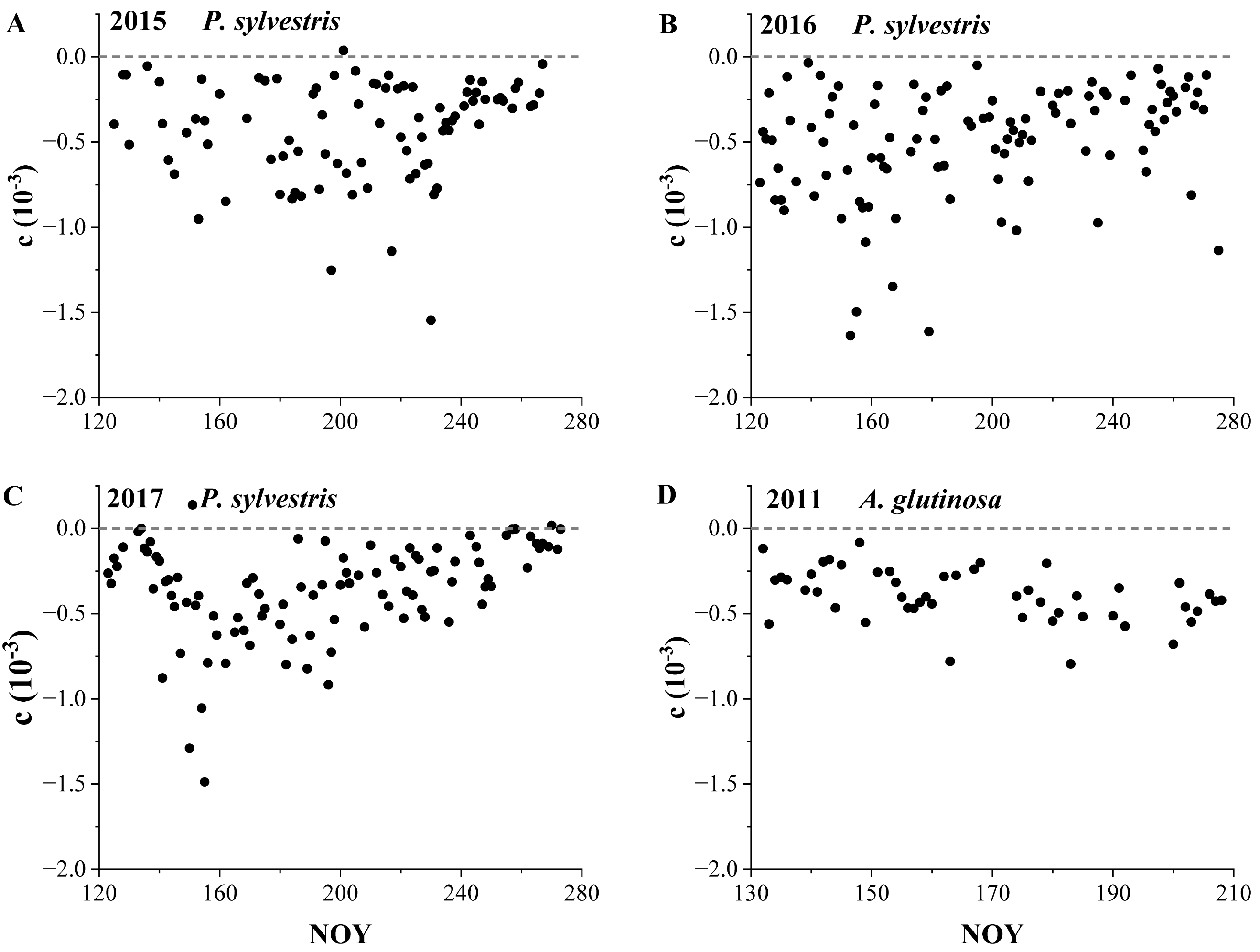

**Fig. S5** Baseline and related sap flux density of *P. sylvestris* and *A. glutinosa*. (A-C) ΔV_max_ and model-obtained baseline (Baseline_new_) (Eqn6). (D-F) Nighttime sap flux density with different baseline (Jn_0_, Jn_new_) (Eq1, Eq7), as well as the sap flux density derived from model (Jn_model_) (Eq5). (G-I) daytime sap flux density with different baseline (Jd_0_, Jd_new_) (Eq1, Eq7). (J-L) Ratio of nighttime sap flux density o daytime with different baseline (Jn_0_/Jd_0_, Jn_new_/Jd_new_). Data resolution: 1 night or day.

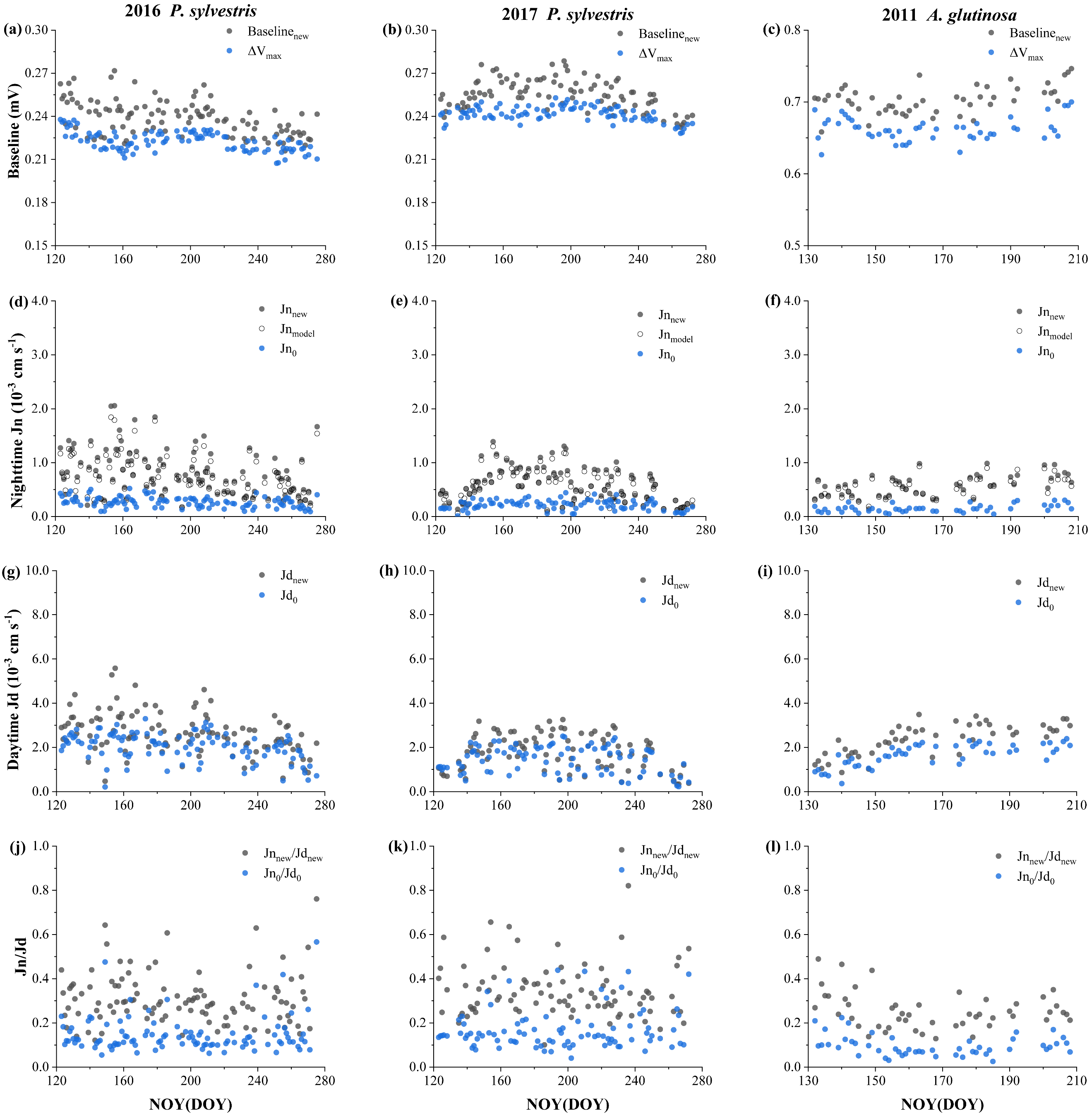

**Fig. S6** Five individual nights per year demonstrating rate of change in diameter (R_Δd_) in relation to sap flux density calculated using ΔV_max_ as baseline (Jn_0_) and using Baseline_new_ as baseline (Jn_new_), and sap flux density derived from model (Jn_model_). Data resolution: 5 min.

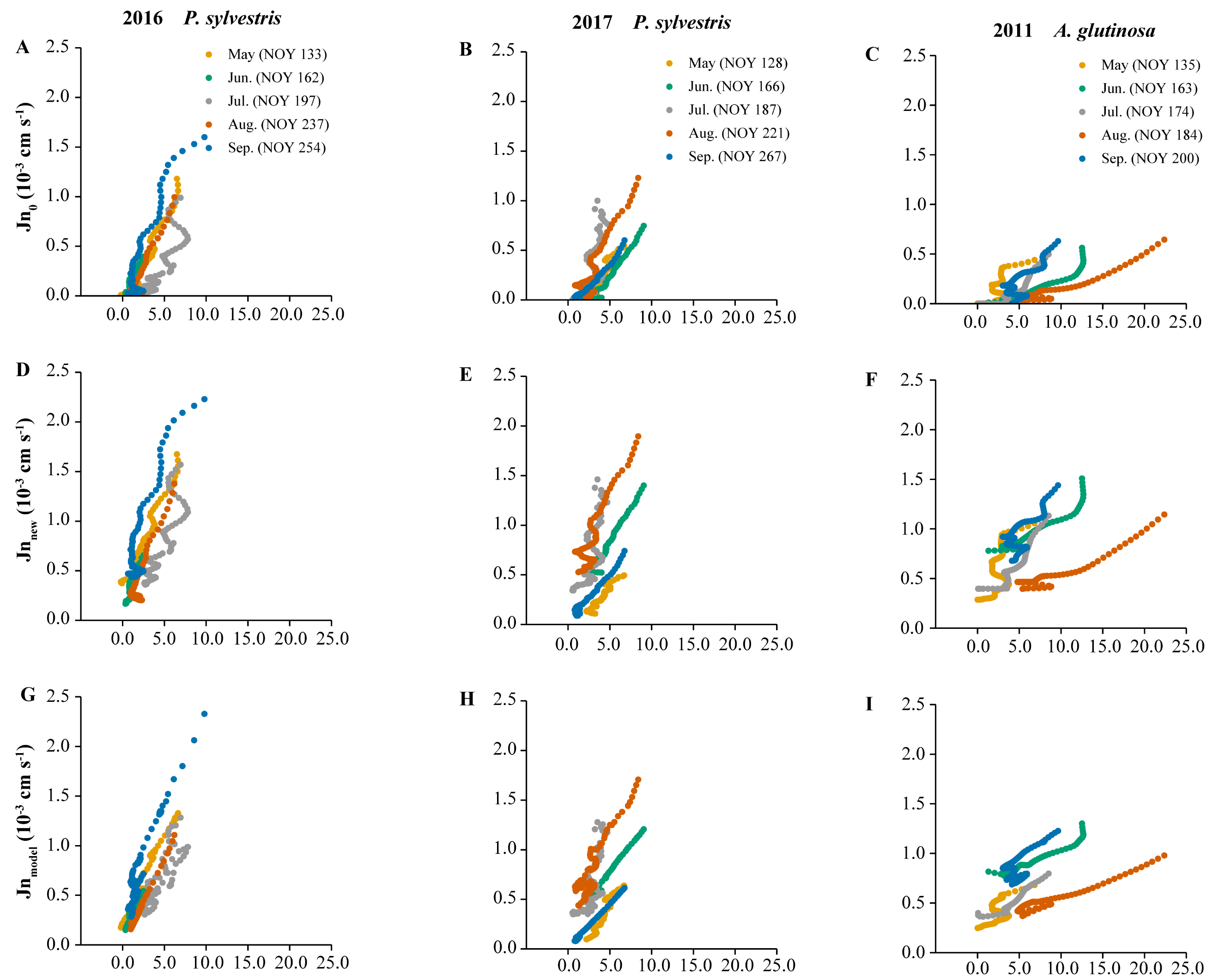

**Fig. S7** Five individual nights per year demonstrating VPD in relation to sap flux density calculated using ΔV_max_ as baseline (Jn_0_) and using Baseline_new_ as baseline (Jn_new_), and sap flux density derived from model (Jn_model_). Data resolution: 5 min.

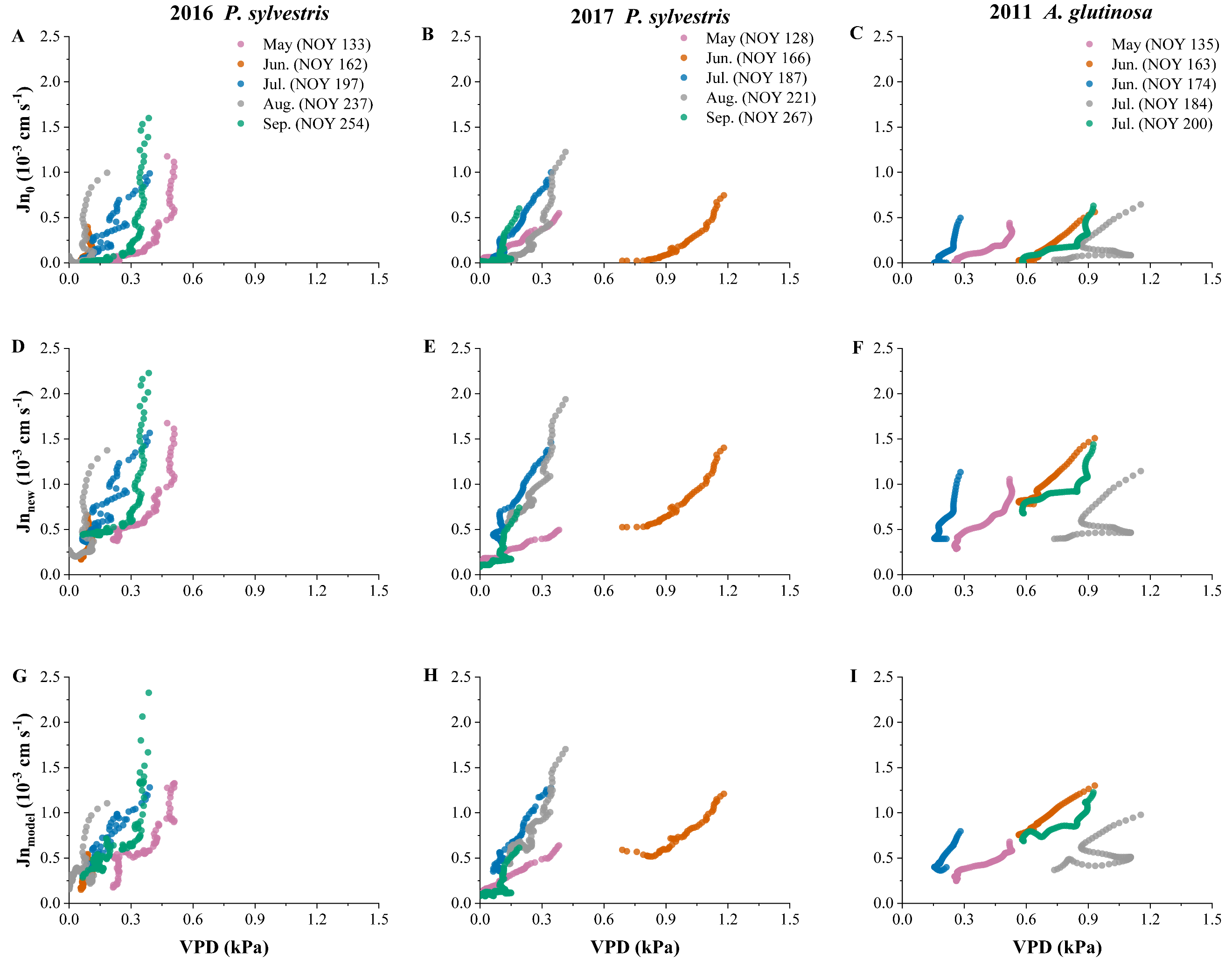

**Fig. S8** Rate of change in diameter (R_Δd_) in relation to sap flux density calculated using ΔV_max_ as baseline (Jn_0_) and baseline derived from model (Jn_new_), and sap flux density derived from model (Jn_model_). Data resolution: 5 min.

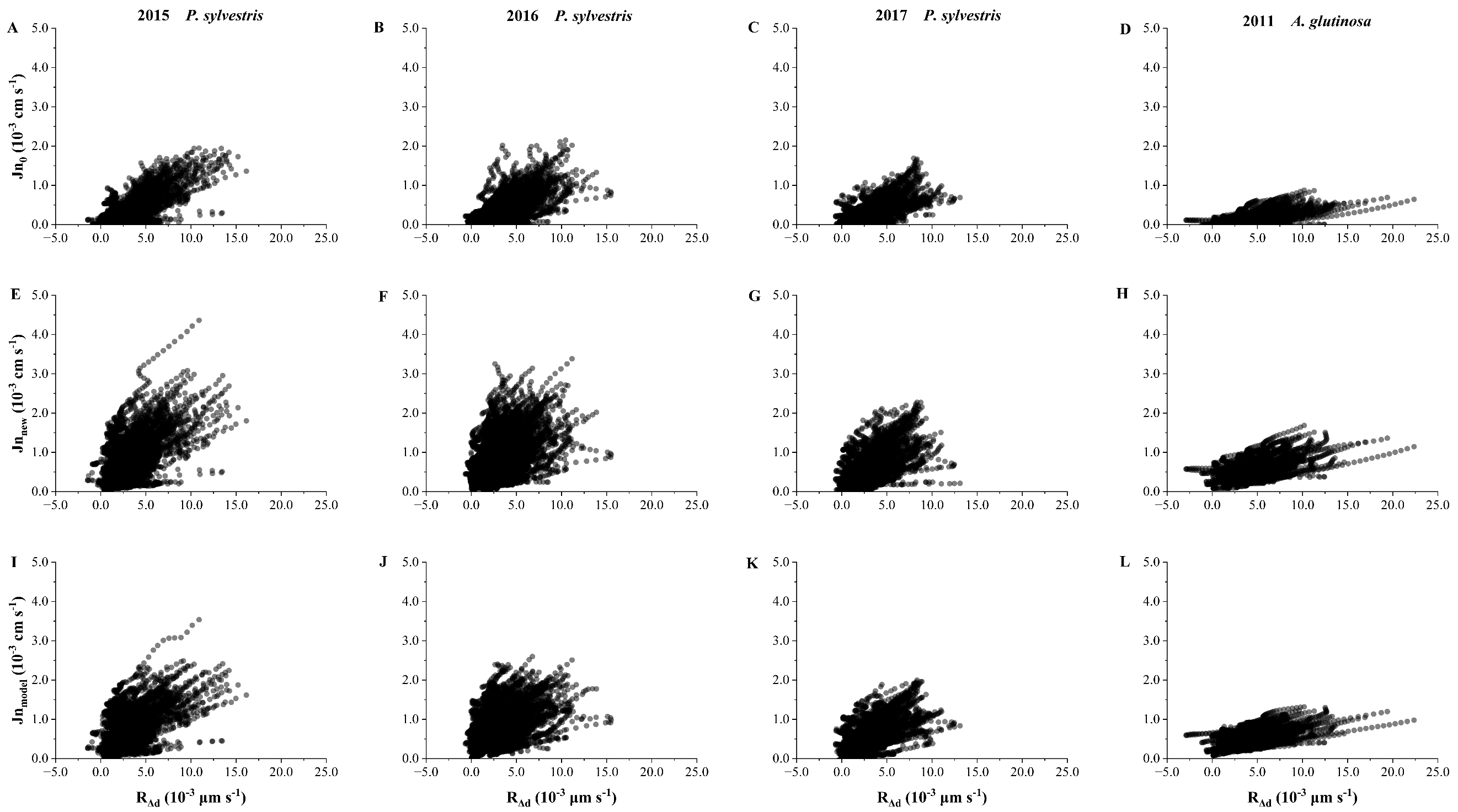

**Fig. S9** VPD in relation to sap flux density calculated using ΔV_max_ as baseline (Jn_0_) and baseline derived from model (Jn_new_), and sap flux density derived from model (Jn_model_). Data resolution: 5 min.

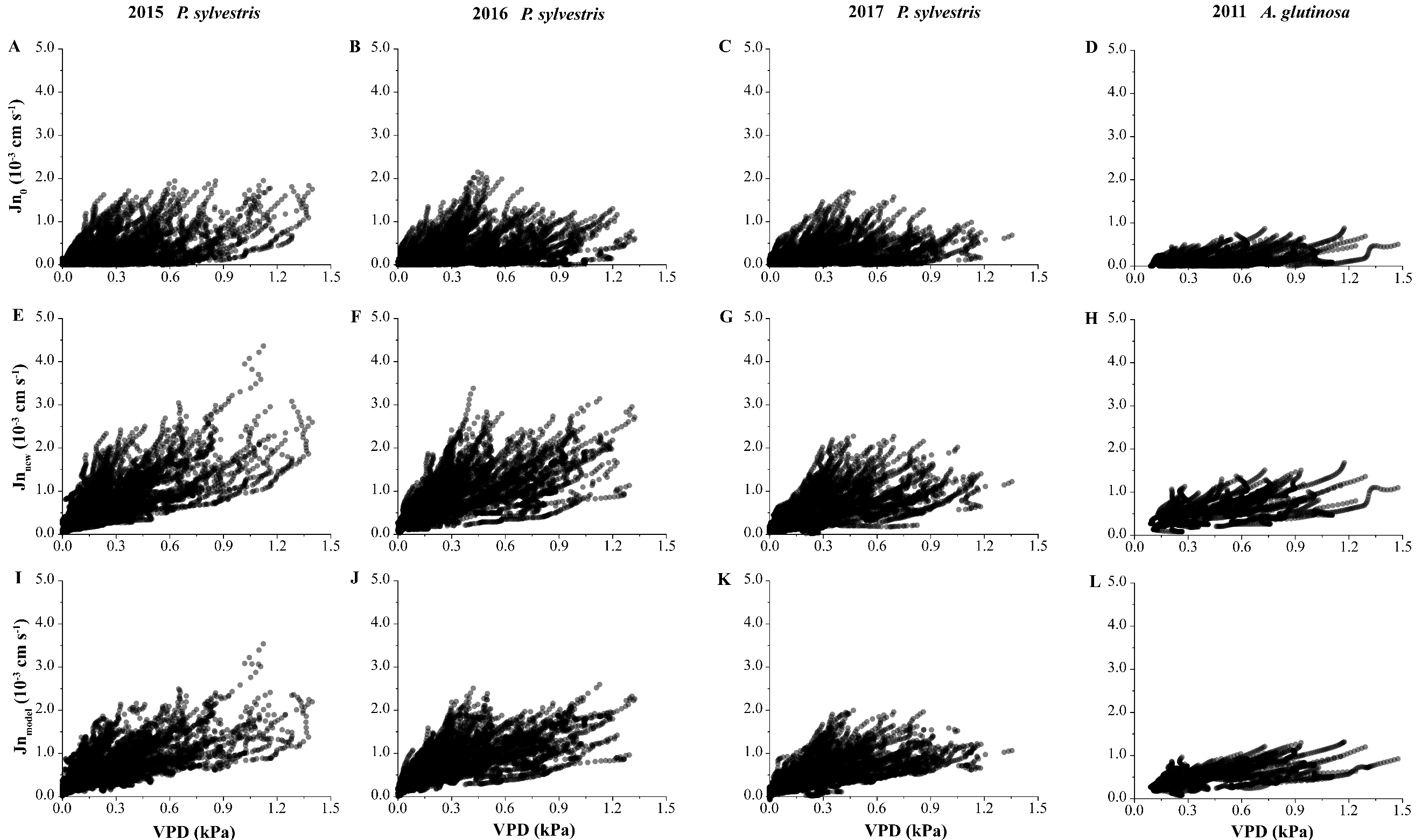

**Fig. S10** Reasons for not being modelled well. Data resolution: 5 min.

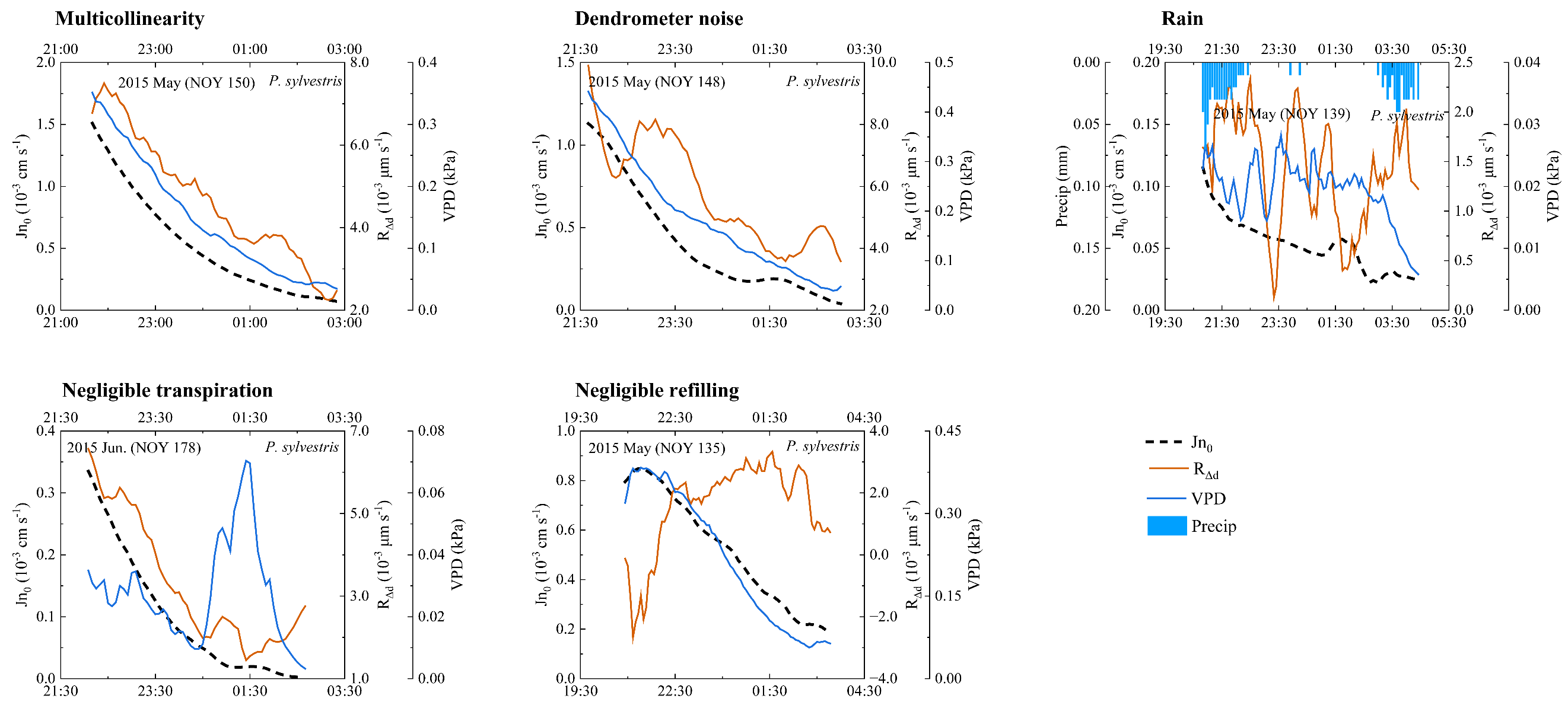

**Fig. S11** R_Adj_^2^ of different models. Data resolution: 1 night.

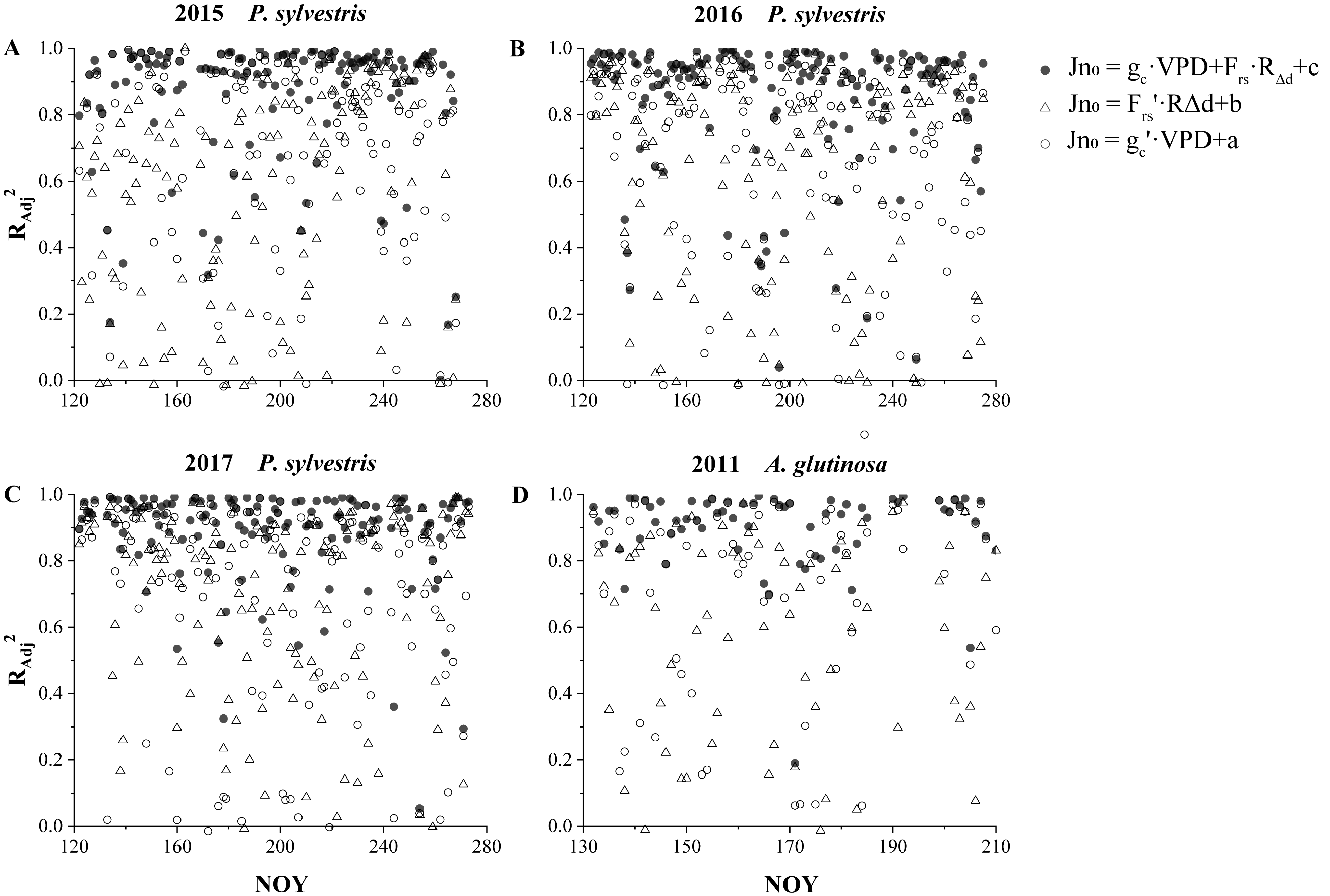

**Fig. S12** Transpiration (T) and refilling (R) flux density. Data resolution: 1 night.

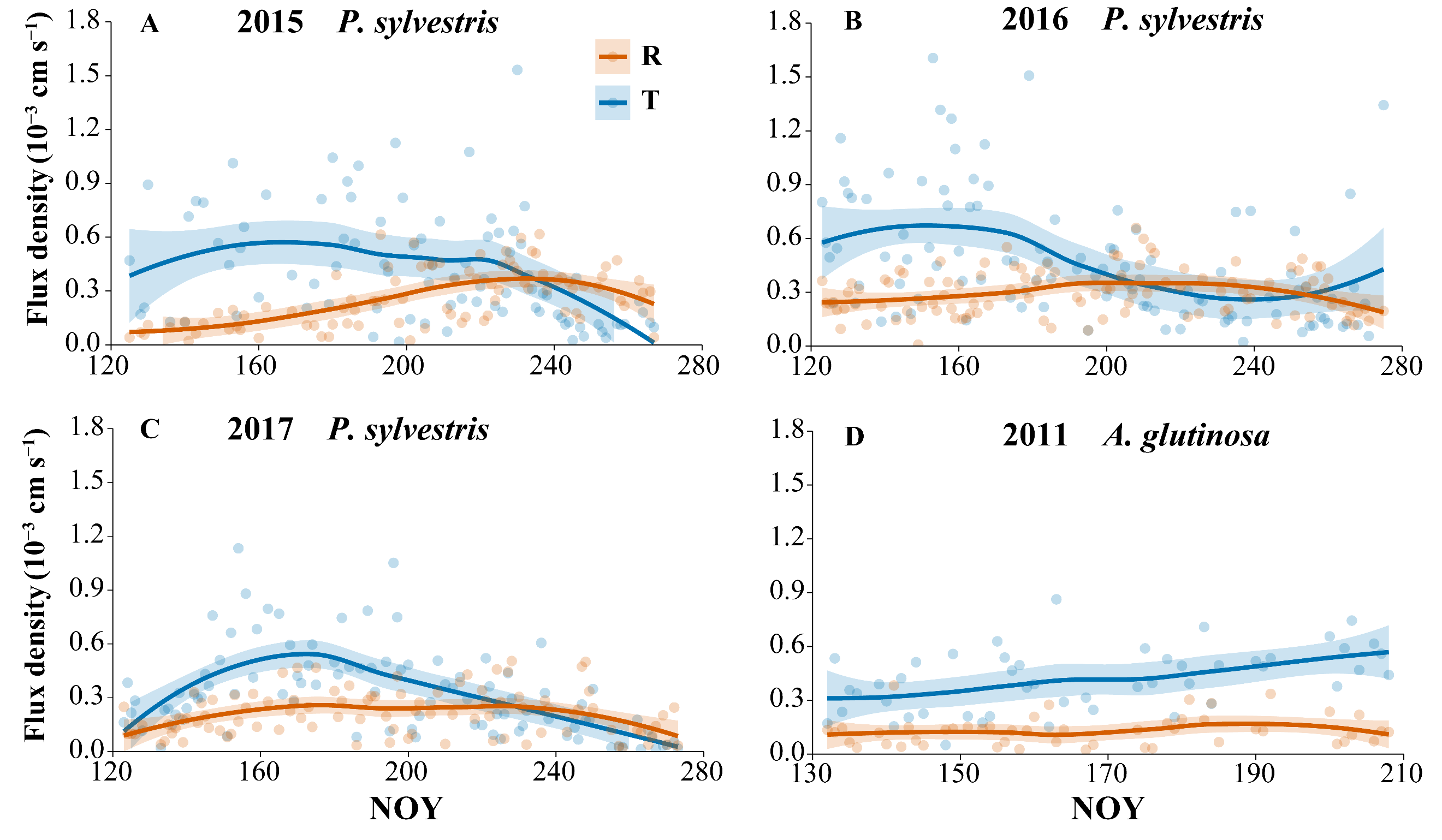

Table S1 Performance distribution of model across nights.

| Performance | | *P. sylvestris* (%) | *A. glutinosa* (%) | Overall (%) |
| --- | --- | --- | --- | --- |
| Modeled nights | | 70.38 | 73.53 | 70.83 |
| Unmodelled nights | Multicollinearity | 3.64 | 7.35 | 4.17 |
|  | Dendrometer noise | 4.13 | 2.94 | 3.96 |
|  | Rain | 14.56 | 10.29 | 13.96 |
|  | Negligible transpiration | 0.24 | 0.00 | 0.21 |
|  | Negligible refilling | 0.49 | 0.00 | 0.42 |
|  | Other reason | 6.55 | 5.88 | 6.46 |

Table S2 R_Adj_^2^ and AIC of different models fitting nighttime sap flux density. Data shown: mean ± SD.

| Model | R_Adj_^2^ | | AIC | |
| --- | --- | --- | --- | --- |
|  | *P. sylvestris* L. | *A. glutinosa* | *P. sylvestris* L. | *A. glutinosa* |
| Jn = g_c_'*VPD+a | 0.70 ± 0.28 bB | 0.72 ± 0.28 bA | -1401.06 ± 459.23 a | -1168.37 ± 189.69 a |
| Jn = F_rs_’*R_Δd_+b | 0.64 ± 0.30 cA | 0.60 ± 0.29 cA | -1389.41 ± 492.44 a | -1120.64 ± 173.75 a |
| Jn = g_c_*VPD+F_rs_*R_Δd_+c | 0.86 ± 0.19 aC | 0.90 ± 0.12 aB | -1487.20 ± 488.75 b | -1232.01 ± 189.25 b |

Different lower-case letters indicate significant differences in R_Adj_^2^ (*P* < 0.01), and different upper-case letters indicate heterogeneity of variance (*P* < 0.05).
